# The DNA methylation landscape of glioblastoma disease progression shows extensive heterogeneity in time and space

**DOI:** 10.1101/173864

**Authors:** Johanna Klughammer, Barbara Kiesel, Thomas Roetzer, Nikolaus Fortelny, Amelie Kuchler, Nathan C. Sheffield, Paul Datlinger, Nadine Peter, Karl-Heinz Nenning, Julia Furtner, Martha Nowosielski, Marco Augustin, Mario Mischkulnig, Thomas Ströbel, Patrizia Moser, Christian F. Freyschlag, Johannes Kerschbaumer, Claudius Thomé, Astrid E. Grams, Günther Stockhammer, Melitta Kitzwoegerer, Stefan Oberndorfer, Franz Marhold, Serge Weis, Johannes Trenkler, Johanna Buchroithner, Josef Pichler, Johannes Haybaeck, Stefanie Krassnig, Kariem Madhy Ali, Gord von Campe, Franz Payer, Camillo Sherif, Julius Preiser, Thomas Hauser, Peter A. Winkler, Waltraud Kleindienst, Franz Würtz, Tanisa Brandner-Kokalj, Martin Stultschnig, Stefan Schweiger, Karin Dieckmann, Matthias Preusser, Georg Langs, Bernhard Baumann, Engelbert Knosp, Georg Widhalm, Christine Marosi, Johannes A. Hainfellner, Adelheid Woehrer, Christoph Bock

## Abstract

Glioblastoma is characterized by widespread genetic and transcriptional heterogeneity, yet little is known about the role of the epigenome in glioblastoma disease progression. Here, we present genome-scale maps of the DNA methylation dynamics in matched primary and recurring glioblastoma tumors, based on a national population registry and a comprehensively annotated clinical cohort. We demonstrate the feasibility of DNA methylation mapping in a large set of routinely collected formalin-fixed paraffin-embedded (FFPE) samples, and we validate bisulfite sequencing as a multi-purpose assay that allowed us to infer a range of different genetic, epigenetic, and transcriptional tumor characteristics. Based on these data, we identified characteristic differences between primary and recurring tumors, links between DNA methylation and the tumor microenvironment, and an association of epigenetic tumor heterogeneity with patient survival. In summary, this study provides a resource for dissecting DNA methylation heterogeneity in genetically diverse and heterogeneous tumors, and it demonstrates the feasibility of integrating epigenomics, radiology, and digital pathology in a representative national cohort, leveraging samples and data collected as part of routine clinical practice.

## Introduction

Glioblastoma is a devastating cancer with a median age at diagnosis of 64 years^1^. Even under the best available care, the median survival is little more than one year, and very few patients live for more than three years^2,3^. Despite intense efforts, limited therapeutic progress has been made over the last decade, and a series of phase III clinical trials with targeted agents have failed to improve overall survival^4-6^.

Glioblastoma shows extensive temporal and spatial heterogeneity, which appears to contribute to therapeutic resistance and inevitable relapse^7-13^. Prior research on tumor heterogeneity in glioblastoma has focused mainly on the genomic and transcriptomic dimensions^7-20^, while the dynamic role of the epigenome in glioblastoma disease progression is much less understood^21^.

Recent data in other cancers have conclusively shown the power of DNA methylation sequencing for analyzing epigenetic heterogeneity. For example, DNA methylation heterogeneity has been linked to clonal progression in prostate cancer^22^, low-grade glioma^23^, esophageal squamous cell carcinoma^24^, and hepatocellular carcinoma^25^; and new measures of DNA methylation heterogeneity such as epi-allele burden, proportion of discordantly methylated reads (PDR), and DNA methylation inferred regulatory activity (MIRA) have been linked to clinical variables in acute myeloid leukemia^26^, chronic lymphatic leukemia^27^, and Ewing sarcoma^28^.

To investigate the contribution of epigenetics to the temporal and spatial heterogeneity of glioblastoma, we performed DNA methylation sequencing on a large cohort of IDH wildtype glioblastoma patients (n = 112) with matched samples from primary and recurring tumors (between 2 and 4 time points per patient), and we also included multiple subregion samples for a subset of these tumors. Importantly, by using an optimized reduced representation bisulfite sequencing (RRBS) protocol^28,29^ we obtained a high success rate for archival formalin-fixed and paraffin-embedded (FFPE) samples, coverage of 3-5 fold more CpGs compared to Infinium microarrays^30-32^, and single-CpG as well as single-allele resolution (in RRBS, each sequencing read captures the DNA methylation status of one or more individual CpGs in one single allele from one single cell).

The presented dataset – comprising 349 RRBS-based DNA methylation profiles, of which 320 were derived from FFPE samples – constitutes the largest cohort of FFPE samples that has yet undergone genome-scale DNA methylation sequencing, and it conclusively shows the technical feasibility of performing large multicenter DNA methylation studies based on routinely collected FFPE material. The RRBS data not only identified epigenetic disease subtypes and quantified epigenetic heterogeneity, but also allowed us to infer transcriptional subtypes, copy number aberration, single nucleotide variants (SNVs), small insertions and deletions (indels), *MGMT* promoter methylation, and G-CIMP/ IDH mutational status. This study thus highlights the power of DNA methylation sequencing in routinely collected clinical FFPE tumor samples.

Linking DNA methylation profiles to magnetic resonance (MR) imaging, tumor morphology, tumor microenvironment, and clinical variables such as patient survival, we obtained a detailed picture of temporal and spatial heterogeneity in glioblastoma. We observed characteristic differences in DNA methylation between primary and recurring tumors, an association with the tumor microenvironment, and a link between tumor microenvironment and previously reported clinically relevant MR imaging-based progression types^33^. DNA methylation was highly predictive of the established glioblastoma transcriptional subtypes, and disease progression-associated loss of DNA methylation in the promoters of Wnt signaling genes was associated with worse prognosis. We also observed across several comparisons that the association with survival was stronger for properties of the recurring tumor than for properties of the primary tumor. In summary, our study provides a comprehensive resource of the DNA methylation dynamics and heterogeneity in glioblastoma, proof-of-concept for DNA methylation sequencing in large FFPE sample sets collected as part of routine diagnostics, and an integrative analysis of DNA methylation with various types of clinical and histopathological information.

## Results

### DNA methylation sequencing in a cohort of matched primary and recurring glioblastoma samples

To investigate the DNA methylation dynamics associated with disease progression in glioblastoma, we established a richly annotated dataset of patients that underwent tumor resection at diagnosis and at least once upon tumor recurrence (Figure 1a). These patients were identified through the population-based Austrian Brain Tumor Registry^34^, and 112 primary glioblastoma patients (wildtype *IDH* status) with tumor samples for at least two time points (primary tumor and first recurrence) were included in the analysis (Supplementary Fig. 1a). Due to the requirement of having undergone at least two tumor resections, the selected patients were on average younger (median age at diagnosis of 58 years) and had longer overall survival (median overall survival of 22.4 months) compared to the unselected population-based cohort (median age at diagnosis 63 years, median overall survival of 8 months) (Figure 1b and Supplementary Table 1).

**Figure 1.**
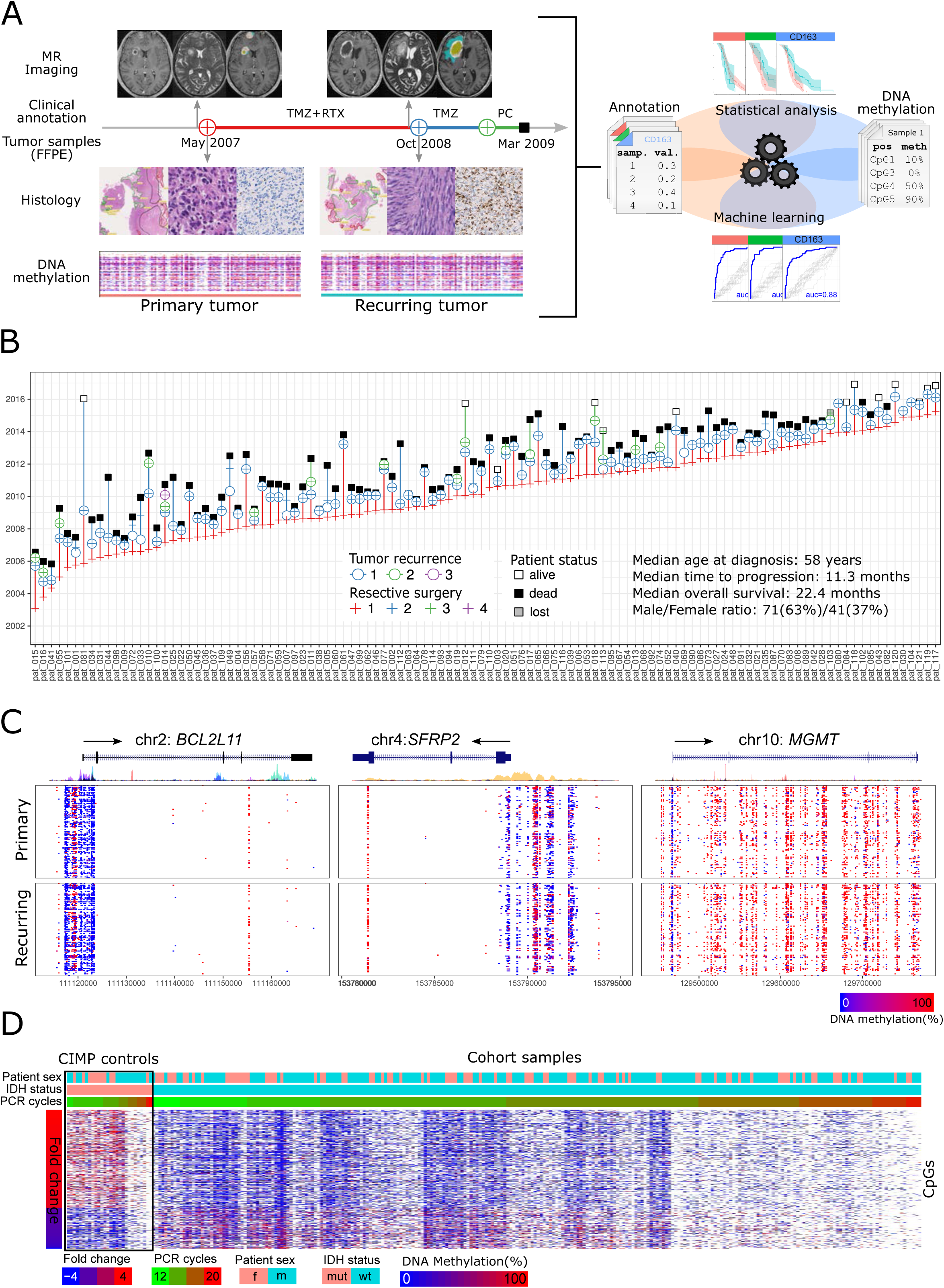
DNA methylation landscape of glioblastoma disease progression. A. Integrative analysis of longitudinal DNA methylation data (RRBS) with matched magnetic resonance (MR) imaging data (morphology, segmentation), clinical annotation data (e.g., treatment, progression, IDH mutation status), and histopathological data (segmentation, morphology, immunohistochemistry) using statistical methods and machine learning. TMZ: Temozolomide; RTX: Radiation therapy; PC: Palliative care.
B. Patient cohort overview summarizing the disease courses of 112 primary glioblastoma patients with IDH-wildtype status, ordered by time of first surgery.
C. DNA methylation profiles for primary and recurring tumors at three relevant gene loci (*BCL2L11*, *SFRP2*, and *MGMT*). Genes and ENCODE histone H3K27ac tracks were obtained from the UCSC Genome Browser.
D. DNA methylation levels at CpGs indicative of the CpG island methylator phenotype (CIMP), shown separately for IDH mutated control samples (which are CIMP-positive) and the IDH wildtype primary glioblastoma samples from the study cohort (which are CIMP-negative). The fold change of DNA methylation levels between IDH mutated and wildtype samples is indicated based on data from a previous study^39^.

For each of these tumor samples, we established genome-scale DNA methylation profiles using reduced representation bisulfite sequencing (RRBS). The RRBS assay provides single-CpG and single-allele resolution with excellent quantitative accuracy even on challenging clinical samples^35,36^, and it has been successfully used to dissect intra-tumor heterogeneity in several cancer types^26-28^. The RRBS-based DNA methylation data were complemented by time-matched MR imaging data as well as quantitative pathology data capturing the morphology, proliferative activity, and tumor microenvironment of the same tumors. All data are available from the Supplementary Website (http://glioblastoma-progression.computational-epigenetics.org/). We systematically integrated these datasets using statistical analysis and machine learning methods (Figure 1a).

DNA methylation profiling based on routinely collected FFPE material was successful for all samples that had adequate tumor cell content, and 95% of the resulting RRBS profiles yielded more than 500,000 covered CpGs (Supplementary Table 2). The median number of covered CpGs in FFPE samples (1,880,675) was lower than for fresh-frozen samples (4,473,349), but higher than for an alternative ethanol-based fixation metho^d37^ (1,005,828) that was tested by one sample-providing center (Supplementary Fig. 1b and Supplementary Table 2). The measured bisulfite conversion rates were highly consistent with expectations: 99% of genomic cytosines outside of CpGs were read as thymines, the mean underconversion rate on unmethylated spike-in controls was 1%, and the mean overconversion rate on methylated spike-in controls was 2% (Supplementary Fig. 1c and Supplementary Table 2). All DNA methylation profiles showed the expected distribution of DNA methylation levels across CpG islands, promoters, and genome-wide tiling regions (Supplementary Fig. 1d), with a tendency toward lower DNA methylation levels in low-quality samples (Supplementary Fig. 1e).

Comparing DNA methylation levels of 5-kilobase tiling regions between primary and recurring tumors, there was a high correlation (r > 0.94) across the genome (Supplementary Fig. 1f). Nevertheless, we observed widespread epigenetic heterogeneity at individual loci (Figure 1c). An example is the promoter of the *MGMT* gene, whose DNA methylation status has been shown to correlate with sensitivity to alkylating chemotherapy^38^. The *MGMT* promoter was unmethylated in the majority of samples (Supplementary Fig. 1g), and patients with a methylated *MGMT* promoter in their recurring tumors had significantly better progression-free survival (PFS) and overall survival (OS) compared to patients with unmethylated *MGMT* promoters (Supplementary Fig. 1h).

To compare our *IDH*-wildtype primary glioblastoma cohort to *IDH*-mutant brain tumor samples, we also performed RRBS on primary and recurring tumors of 13 *IDH*-mutant oligodendroglioma/astrocytoma/glioblastoma patients from the same population. The DNA methylation profiles of these tumors showed a characteristic CpG island methylator phenotype (CIMP) as expected from previous observations^39^ (Figure 1d), which provides further validation of the accuracy and robustness of our DNA methylation profiling on FFPE material.

### Inference of genomic information from the RRBS data

All tumor samples were obtained via a national brain tumor registry and reflect routine clinical care in Austria, which currently does not include whole genome or whole exome sequencing. We therefore evaluated whether certain types of genomic information could be inferred directly from the RRBS data.

First, we reconstructed genome-wide maps of copy number aberrations (CNAs) from the RRBS data using the CopywriteR algorithm^40^. We detected various CNAs previously described in glioblastoma, including amplifications of the *EGFR* locus and deletions of chromosome 10 (Supplementary Fig 2a). In particular, we observed that 10q deletions in the recurring tumors (which affect the *MGMT* gene and have been shown to correlate with sensitivity to alkylating chemotherapy^41^) were associated with longer survival (Supplementary Fig. 2b). Based on the CNA data, we also verified that none of our primary glioblastoma samples harbored the 1p19q co-deletion, thereby excluding the presence of any misclassified cases of anaplastic oligodendroglioma (Supplementary Fig. 2c).

Second, we inferred single nucleotide variants (SNVs) and small insertions/deletions (indels) from the RRBS data using the Bis-SNP algorithm^42^. Although confident SNV and indel detection in RRBS data is limited to a relatively small subset of the genome, it allowed us to confirm that none of our samples displayed a hypermutator phenotype and that there was no strong trend toward higher mutational rates in primary versus recurring tumors (Supplementary Fig. 2d). Furthermore, among the variants with high predicted impact on protein expression we found multiple genes with known relevance in glioblastoma (Supplementary Fig 2e).

### Prediction of transcriptional subtypes in glioblastoma based on DNA methylation

Recent research defined three transcriptional subtypes of glioblastoma (classical, mesenchymal, and proneural)^43^, while a previously described fourth (neural) subtype has been described as an artifact of contaminating non-tumor tissue. Because RNA sequencing of FFPE material is challenging and often infeasible, we tested whether these transcriptional subtypes can be inferred from DNA methylation data (Figure 2a). Machine learning classifiers using L2-regularized logistic regression were trained and evaluated on matched DNA methylation and transcriptional subtype data^44,45^ from The Cancer Genome Atlas (http://cancergenome.nih.gov/). Based on these data, we obtained good prediction accuracies with cross-validated receiver operating characteristic (ROC) area under curve (AUC) values above 0.8 for most samples (Supplementary Fig 3a).

**Figure 2.**
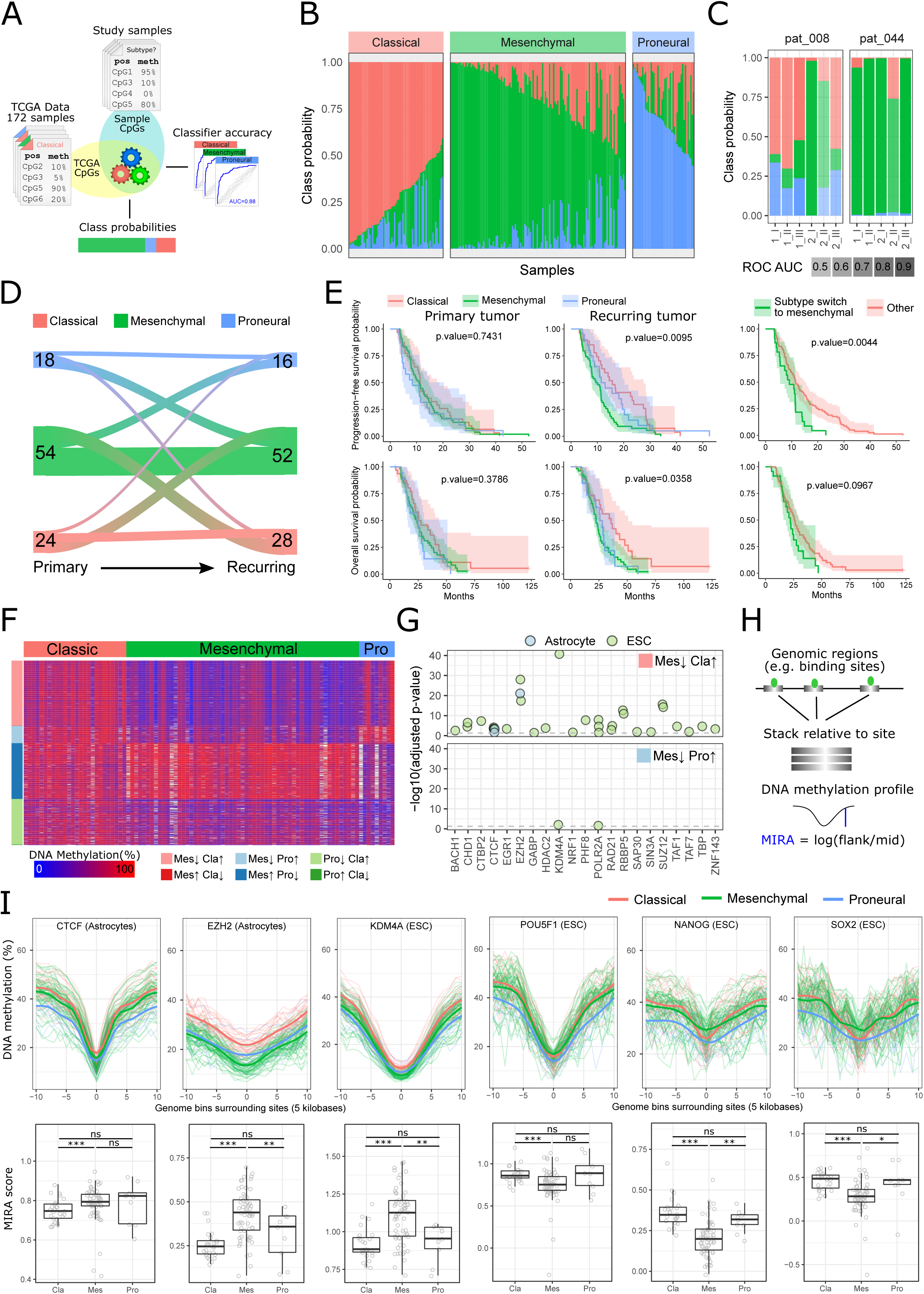
Glioblastoma transcriptional subtypes inferred from DNA methylation. A. Overview of the machine learning approach for classifying tumor samples by their transcriptional subtypes using DNA methylation data. Classifiers were trained on DNA methylation data (Infinium 27k assay) of TCGA glioblastoma samples with known transcriptional subtype, using only CpGs shared by RRBS. All classifiers were evaluated by tenfold cross-validation on the TCGA samples and then applied to the RRBS profiles, predicting class probabilities that indicate the relative contribution of each transcriptional subtype.
B. Transcriptional subtype heterogeneity within cohort samples, as indicated by class probabilities of the subtype classifier. Samples are grouped and ordered by their dominant subtype.
C. Distribution of class probabilities across different regions of the same tumor (indicated by Roman numbers) and across different surgeries (indicated by Arabic numbers) for two patients with multisector samples.
D. Riverplot depicting transitions in the predicted transcriptional subtype between primary and recurring tumors. The number of samples in each state is indicated. Only patients whose primary and recurring tumors were classified with high accuracy (ROC AUC > 0.8) were included in this analysis.
E. Kaplan-Meier plots displaying progression-free survival and overall survival probabilities over time for patients stratified by predicted transcriptional subtypes (left) and switching from a non-mesenchymal to mesenchymal subtype during disease progression (right). Only tumor samples that were classified with high accuracy (ROC AUC > 0.8) were included in this analysis.
F. Heatmap displaying the DNA methylation levels of the most differential CpGs between the three transcriptional subtypes. Only tumor samples that were classified with high class probabilities (>0.8) were included in this analysis.
G. LOLA region-set enrichment analysis of differentially methylated CpGs between the different transcriptional subtypes (binned into 1-kilobase tiling regions). Adjusted p-values (Benjamini & Yekutieli method) are displayed for all significantly (adjusted p-value < 0.05) enriched region sets (binding sites, x-axis) measured in astrocytes or embryonic stem cells (ESCs).
H. Schematic depicting the calculation of ‘DNA methylation inferred regulatory activity’ (MIRA) scores. DNA methylation profiles are combined across centered genomic regions of interest (e.g., transcription factor binding sites) for each sample and each region set. The MIRA score is then calculated as the ratio of DNA methylation levels at the flank to DNA methylation levels at the center (binding site) of the combined DNA methylation profile. High MIRA scores therefore reflect local demethylation at the binding site, which indicates high regulatory activity of the respective factor.
I. DNA methylation profiles (upper row) and corresponding MIRA scores (lower row) for three region sets enriched in CpGs that are hypomethylated in the mesenchymal subtype (CTCF binding in astrocytes, EZH2 binding in astrocytes, and KDMA binding in ESCs) as well as three region sets of key regulators of pluripotency measured in ESCs (POUF1, NANOG, SOX2). The significance of differences between the three transcriptional subtypes was assessed using a two-sided Wilcoxon rank sum test: ^*^ p-value < 0.05, ^**^ p-value < 0.01, ^***^ p-value < 0.001, ns: not significant.

Applying the trained classifiers to our DNA methylation dataset, we assigned class probabilities to each tumor sample. While these percentage values primarily reflect the confidence with which each sample is assigned to each of the transcriptional subtypes, here we used them as an indicator of the relative contribution of each of the three subtypes to individual tumor samples, thus providing an initial assessment of intra-tumor heterogeneity. We found that all three transcriptional subtypes were common in our cohort of *IDH* wildtype primary glioblastoma (Figure 2b) – in contrast to the *IDH* mutated tumors, which were almost always assigned to the proneural subtype (Supplementary Fig 3b).

Predicted transcriptional subtypes were heterogeneous both in space and in time. Most individual tumor samples showed signatures of more than one transcriptional subtype (Figure 2b), which is consistent with recent single-cell RNA-seq data that identified similar heterogeneity within individual samples^11^. Moreover, five out of six patients with multi-sector samples displayed at least two transcriptional subtypes (Figure 2c and Supplementary Fig. 3c-e), and about half of the patients showed different predominant transcriptional subtypes between the primary and recurring tumors (Figure 2d). Predicted transcriptional subtypes in the recurring tumor (but not in the primary tumor) were also associated with patient survival (Figure 2e), the mesenchymal subtype being associated with the worst prognosis and the classical subtype with the best prognosis. Finally, patients whose tumors switched to the mesenchymal subtype displayed the worst PFS and OS (Figure 2e).

To investigate the epigenetic differences between the three transcriptional subtypes, we compared the DNA methylation profiles of those tumors that were most confidently assigned to one specific subtype (class probability > 0.8) with each other (Figure 2f). Performing region set analysis for the differentially methylated CpGs using LOLA^46^, we identified a moderate enrichment of chromatin protein binding sites among regions hypomethylated in the mesenchymal subtype, including EZH2, KDM4A, RBBP5, and SUZ12 (Figure 2g).

We also calculated ‘DNA methylation inferred regulatory activity’ (MIRA) scores^28^ for each individual tumor sample, where high scores indicate strong local depletion of DNA methylation at specific transcription factor binding sites (Figure 2h). We observed significantly higher MIRA scores (corresponding to deeper DNA methylation dips and increased regulatory activity) for CTCF, EZH2, and KDM4A in the mesenchymal subtype (Figure 2i). In contrast, MIRA scores for key regulators of pluripotency (NANOG, SOX2, POU5F1) were reduced in the mesenchymal subtype (Figure 2i). To corroborate this observation, we plotted MIRA scores for EZH2 and NANOG against class probabilities for the different transcriptional subtypes across all samples (not only those with high class probability), and we indeed observed highly significant correlations with mesenchymal class probabilities of 0.49 (EZH2) and -0.48 (NANOG) (Supplementary Fig. 3f).

### Linking DNA methylation differences to changes in the tumor microenvironment

To test whether the DNA methylation data captures relevant aspects of the tumor microenvironment, we quantified the abundance of various types of immune cells in the primary and recurring tumors. Specifically, we performed single-plex stainings for markers that identified different immune cell types (CD3, CD8, CD80, CD68) including anti-inflammatory (CD163, FOXP3) and memory T-cell (CD45ro) subpopulations. Similarly, cell proliferation was measured by staining for the cell proliferation marker Ki67 (MIB1) followed by quantification of the relative abundance of MIB1 positive cells. We observed significant differences in the immune cell count between the three transcriptional subtypes (Figure 3a-b), with the highest number of immune cells found in tumors of the mesenchymal subtype. The increased level of immune cell infiltration was accompanied by larger necrotic areas, fewer vital tumor areas, lower tumor cell proliferation, and lower cell density in tumor tissues (Supplementary Fig. 4a-b). Moreover, high levels of CD68 positive cells (macrophages of all types) were associated with poor prognosis in recurring tumors; and high levels of CD163 positive cells (anti-inflammatory, tumor-propagating M2 macrophages) were associated with poor prognosis in both primary and recurring tumors (Figure 3c).

**Figure 3.**
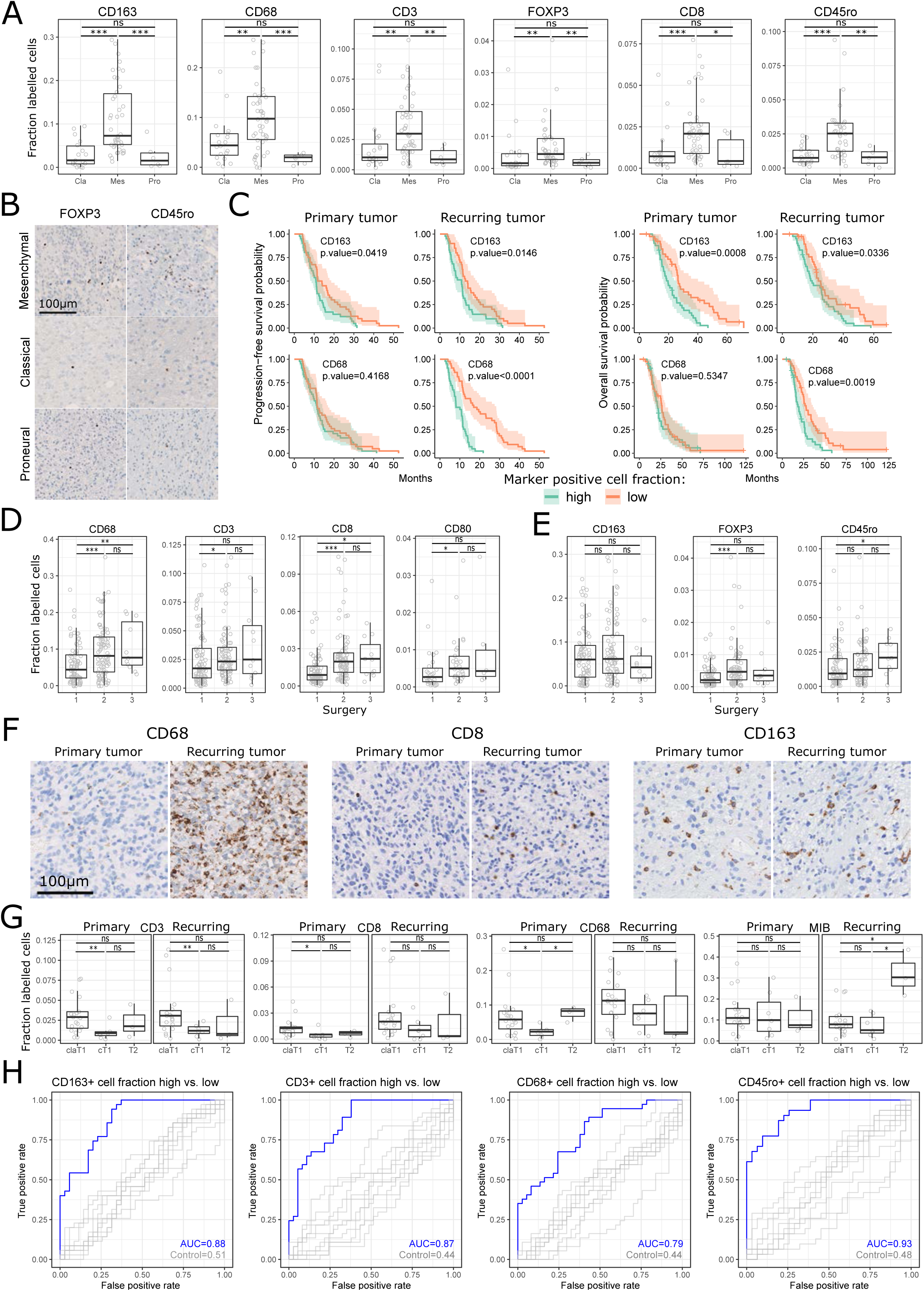
DNA methylation and the tumor microenvironment. A Comparison of tumor-infiltrating immune cell levels between different transcriptional subtypes as measured by quantitative immunohistochemistry for the indicated marker proteins.
B Immunohistochemical stainings for FOXP3 and CD45ro in selected samples assigned to each of the three transcriptional subtypes.
C Kaplan-Meier plots displaying progression-free survival and overall survival probabilities over time for patients stratified according to the level of CD163-positive and CD68-positive immune cell infiltration in their primary and recurring tumors.
D, E Differences in the relative proportion of tumor-infiltrating pro-inflammatory (D) and anti-inflammatory or neutral (E) immune cells between tumor samples originating from the patients’ first surgery (primary tumor), second surgery (recurring tumor), or third surgery.
F Comparative immunohistochemical stainings between primary and recurring tumors for three selected markers (CD68, CD8, CD163).
G Comparison of the levels of tumor-infiltrating immune cells (cells positive for CD3, CD8, or CD68) and proliferating cells (MIB-positive cells) between the different progression types based on magnetic resonance (MR) imaging: Classic T1 (claT1), cT1 relapse / flare-up (cT1), and T2 diffuse (T2). Primary (left panel) and recurring (right panel) tumors were analyzed separately.
H ROC curves for the DNA methylation based prediction of immune cell infiltration levels, as determined by leave-one-out cross-validation. ROC curves and the ROC area under curve (AUC) are indicated for the actual prediction (blue) and for background predictions with randomly shuffled labels (grey). All significance tests comparing groups of samples in this figure were performed using a two-sided Wilcoxon rank sum test: ^*^ p-value < 0.05, ^**^ p-value < 0.01, ^***^ p-value < 0.001, ns: not significant.

Comparing immune cell infiltration between primary and recurring tumors of the same patients, we observed significantly increased levels of inflammatory infiltrates upon recurrence, while the infiltration levels of antiinflammatory macrophages and memory T-cells did not change significantly (Figure 3d-f). The increase in inflammation in the recurring tumors was accompanied by a decrease in necrotic volume and contrast-enhancing (active) tumor mass, but an increase in edema according to matched diagnostic MR imaging data (Supplementary Fig 4c-d). Patients with less necrotic or contrast-enhancing (active) tumor mass upon recurrence presented with a more favorable clinical outcome (Supplementary Fig. 4e).

When we stratified patients according to prognostically relevant MR imaging-based progression types (i.e., classic T1, cT1 relapse / flare-up, and T2 diffuse)^33^ (Supplementary Fig. 5a), we found that primary and recurring tumors from patients displaying the “cT1 relapse / flare-up” subtype had lower infiltration of pro-inflammatory immune cell types (CD3, CD8, CD68) and a lower fraction of proliferating cells (MIB) (Figure 3g). In concordance with previous work^33^, cT1 relapse / flare-up patients displayed a slightly better survival compared to the other two progression types (Supplementary Fig. 5b).

Several of these characteristics of the tumor microenvironment could be predicted from the DNA methylation data using machine learning methods (Supplementary Fig. 5c-d). Differentiating between tumors with a high and low level of immune cell infiltration, we observed a high cross-validated prediction performance for CD163 (ROC AUC = 0.88), CD68 (0.79), CD45ro (0.93), CD3 (0.87) and CD8 (0.79) (Figure 3h; Supplementary Fig. 5e). For those immune cell types with high cross-validation accuracy, DNA methylation levels at the most predictive genomic tiling regions accurately grouped the samples by their immune cell infiltration levels in a hierarchical clustering analysis (Supplementary Fig 5f). Extending recent research that has demonstrated the feasibility of inferring immune cell infiltration from RNA expression profiles^20,47^, our data support that DNA methylation data can be used in similar ways.

### Linking DNA methylation differences to tumor cell-intrinsic properties

To investigate the relationship between DNA methylation and tumor cell characteristics such as proliferation and nuclear morphology, we performed detailed histopathological analysis for the majority of tumor samples.

The percentage of proliferating (MIB positive) cells showed no consistent changes between primary and recurring tumors (Figure 4a). Nevertheless, high proliferation in the recurring tumors (but not in the primary tumors) was significantly associated with increased PFS (Figure 4b). DNA methylation patterns discriminated with high accuracy between tumors characterized by high versus low proliferation rates (ROC AUC = 0.89) (Figure 4c). This was not due to differences in mean DNA methylation levels (Figure 4d-e). Rather, highly proliferating tumors showed intermediate DNA methylation levels at discriminatory regions, while low proliferating tumors showed more extreme methylation patterns (Figure 4e).

**Figure 4.**
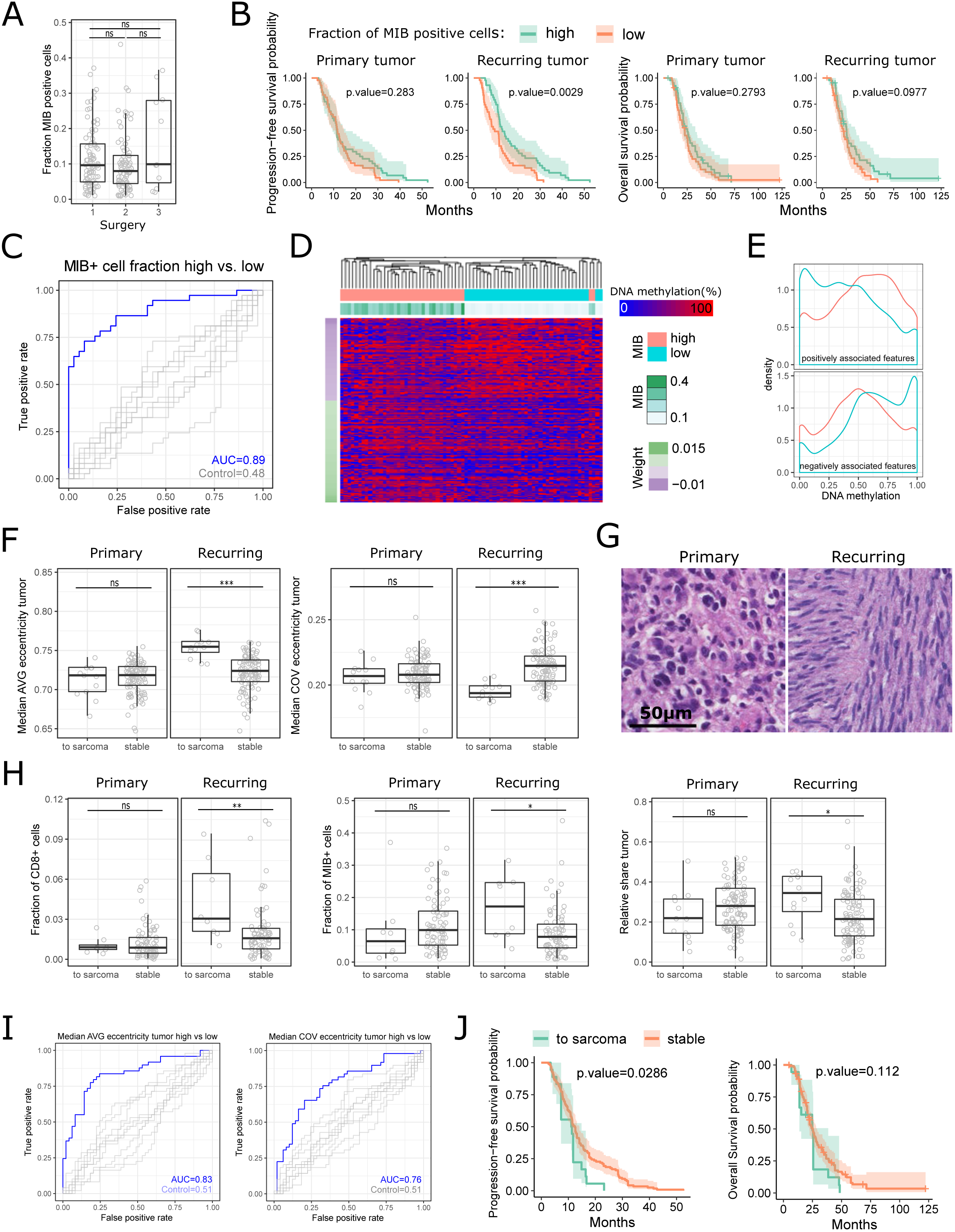
DNA methylation and histopathological tumor characteristics. A. Comparison of the fraction of proliferating (MIB-positive) cells between first surgery (primary tumor), second surgery (recurring tumor), and third surgery.
B. Kaplan-Meier plots displaying progression-free survival and overall survival probabilities over time for patients stratified according to the fraction of proliferating (MIB-positive) cells in the primary and recurring tumor.
C. ROC curves for the DNA methylation based prediction of levels of proliferating (MIB positive) cells, as determined by leave-one-out cross-validation.
D. Hierarchical clustering based on column-scaled DNA methylation levels of the most predictive genomic regions from the classifier predicting the fraction of proliferating (MIB-positive) cells (5-kilobase tiling regions).
E. Distribution of DNA methylation levels of the most predictive genomic regions from the classifier predicting the fraction of proliferating (MIB-positive) cells, displayed separately for samples with high (red) or low (blue) fractions and for positively (top) or negatively (bottom) associated features as identified by the classifier.
F. Comparison of the average nuclear eccentricity (AVG) and its coefficient of variation (COV) between tumors that shift to a sarcoma-like phenotype during disease progression and those that retain a stable histological phenotype.
G. Hematoxylin and eosin stains of matched primary and recurring tumors, illustrating the morphological changes observed when tumors shift to a sarcoma-like phenotype during disease progression.
H. Comparison of additional tumor properties between tumors that shift to a sarcoma-like phenotype during disease progression and those that retain a stable histological phenotype.
I. ROC curves for the DNA methylation based prediction of average nuclear eccentricity and its coefficient of variation.
J. Kaplan-Meier plots displaying progression-free survival and overall survival probabilities over time for patients stratified according to whether their tumors shift to a sarcoma-like phenotype during disease progression (green) or retain a stable histological phenotype (orange). All significance tests comparing groups of samples in this figure were performed using a two-sided Wilcoxon rank sum test: ^*^ p-value < 0.05, ^**^ p-value < 0.01, ^***^ p-value < 0.001, ns: not significant.

Nuclear morphology of tumor cells was measured by the size and eccentricity (a measure of elongated shape) of the tumor cell nuclei, and by the variability of the two parameters. None of these measures were significantly different between primary and recurring tumors. However, tumors that shifted to a sarcoma-like phenotype (i.e., secondary gliosarcoma) upon recurrence had a significant increase in nuclear eccentricity and a decrease in its variability (Figure 4f-g), accompanied by an increase in CD8 immune cell infiltration, proliferative rates (MIB+ cells), and relative tumor mass in the recurring tumor (Figure 4h). DNA methylation patterns predicted nuclear eccentricity (ROC AUC = 0.83) and its variability (0.76) (Figure 4i), and patients with shape-shifting tumors (i.e., classic to sarcoma) displayed significantly shorter PFS and a trend towards reduced OS (Figure 4j), which might be explained by increased tissue infiltration of the spindle-shaped cells.

### DNA methylation heterogeneity and dynamics between primary and recurring tumors

To quantify epigenetic tumor heterogeneity in glioblastoma progression, we used two complementary approaches (Figure 5a). Sub-clonal heterogeneity was measured by epi-allele entropy^48^, and stochastic DNA methylation erosion was measured by the proportion of discordant reads (PDR)^27^. Both measures identified extensive heterogeneity between patients (Supplementary Fig. 6a-b) but no strong trend between primary and recurring tumors (Figure 5b).

**Figure 5.**
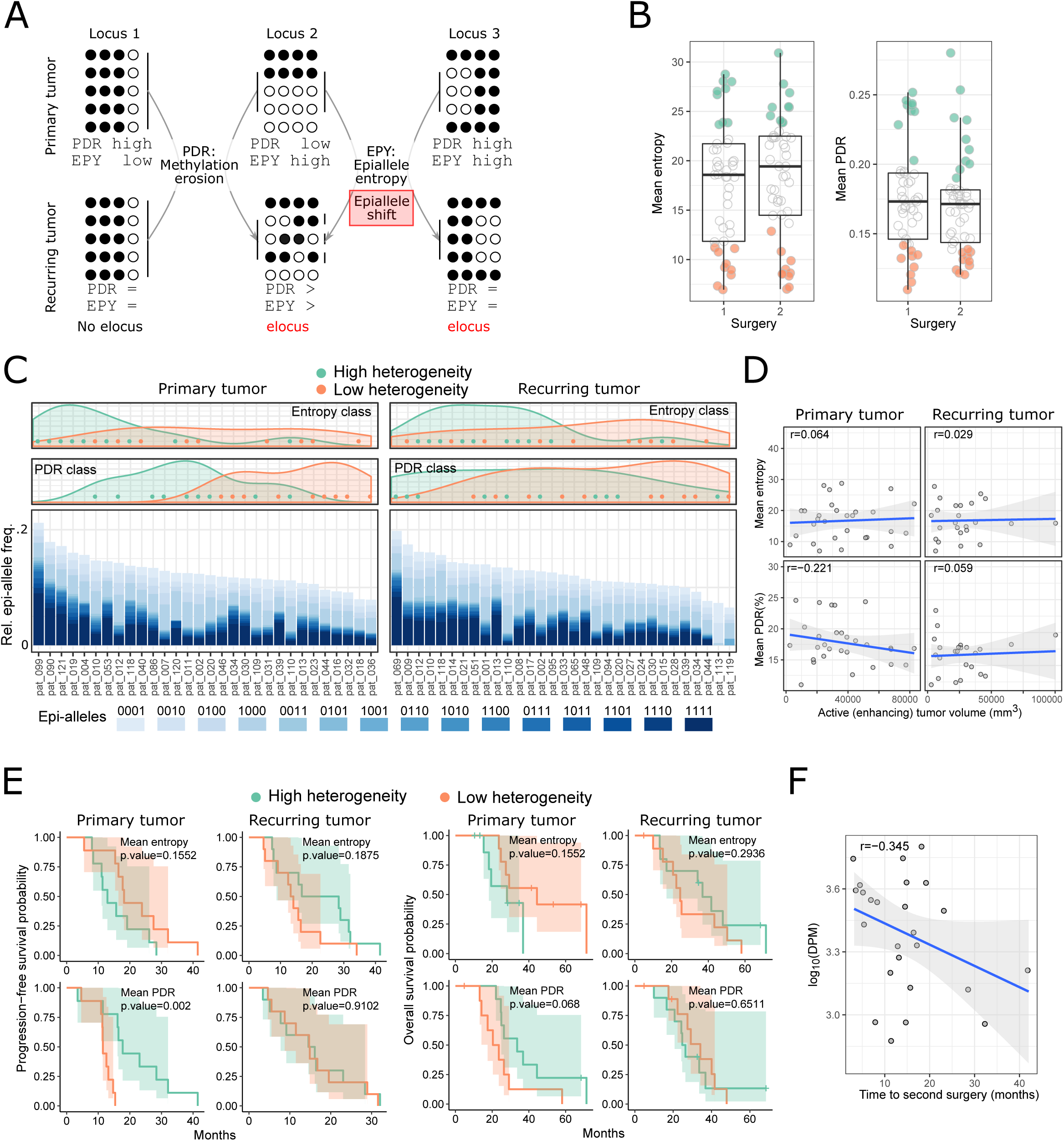
DNA methylation heterogeneity in glioblastoma disease progression. A. Illustration of epi-allele entropy (EPY) and proportion of discordant reads (PDR) as two complementary measures of epigenetic tumor heterogeneity. Individual loci can have high values for one but not for the other measure (locus 1 and 2), or the measures can agree with each other (locus 3). Loci that undergo extensive changes in their epi-allele composition (locus 2 and 3) have been termed eloci^48^. ‘=’: no change in heterogeneity between primary and recurring tumor; ‘>’ increased heterogeneity in the recurring tumor.
B. Comparison of mean sample-wise PDR and epi-allele entropy values between first surgery (primary tumor) and second surgery (recurring tumor). The samples with the 20% highest and lowest heterogeneity values are color-coded and form the basis for the analyses presented in panels C to E.
C. Relative epi-allele frequencies in promoter regions for each of the color-coded samples from panel B (bottom) as well as the distribution of high and low heterogeneity samples along the gradient defined by their relative epi-allele composition (top). For clearer visualization, the “0000” majority epi-allele with a frequency of 70% to 80% is not displayed. 0: unmethylated, 1: methylated
D. Correlation between intra-tumor heterogeneity and enhancing (active) tumor mass as determined by MR imaging. r: Pearson correlation.
E. Kaplan-Meier plots displaying progression-free survival and overall survival probabilities over time for patients stratified according to their PDR and epi-allele entropy values.
F. Correlation between the number of differentially methylated promoters during progression per million assessed promoters (DPM) and the time between first and second surgery. r: Pearson correlation.

Comparing the epi-allele composition of the 20% most heterogeneous and the 20% least heterogeneous samples (Figure 5b), we found extensive variability between patients and over time (Figure 5c). Samples with high epi-allele entropy and high diversity in their epi-allele composition also showed high PDR values, indicating that these samples were characterized by broadly increased epigenetic heterogeneity.

The observed differences in epigenetic heterogeneity were not a side effect of different tumor sizes, as there was little to no correlation between the two measures of tumor heterogeneity on the one hand and the tumor size as measured by MR imaging on the other hand (Figure 5d). In contrast, we did observe a significant association between PDR values and clinical outcome specifically in the primary tumors (Figure 5e and Supplementary Fig. 6c), and we also found that a longer time span between first and second surgery was weakly associated with fewer differentially methylated regions (Figure 5f) but not with the extent of epi-allelic shifting (Supplementary Fig. 6d).

To further dissect the temporal dimension of DNA methylation heterogeneity in glioblastoma, we performed differential DNA methylation analysis on all matched pairs of primary and first recurring tumors. Focusing on gene promoters, we observed high correlation of DNA methylation levels (r = 0.86) and a small number of promoters with strong differential methylation in multiple patients (Figure 6a). Most of these promoters showed consistent trends toward either gain or loss of DNA methylation upon tumor recurrence, although for one gene with known involvement in brain cancer (*OTX2*) we observed a progression-associated gain of DNA methylation in some patients and a loss in others (Figure 6b upper panel). When we classified the patients into those that followed the cohort-level trend in differential DNA methylation (trend patients) and those that did not (anti-trend patients) (Figure 6b, lower panel and Figure 6c), the trend patients showed worse prognosis (Figure 6e), suggesting that some of the observed differences may contribute to disease progression.

**Figure 6.**
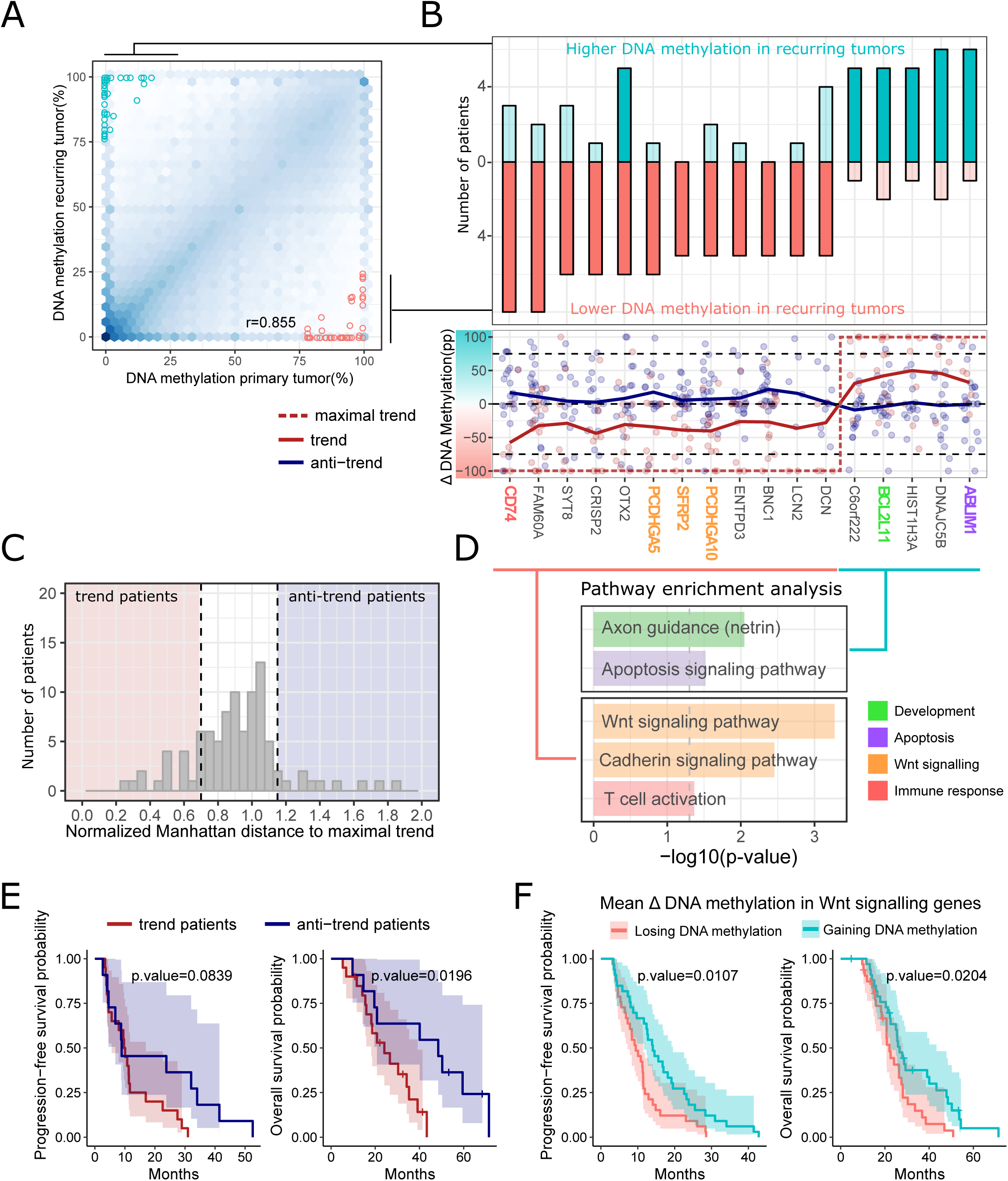
DNA methylation differences between primary and recurring tumors. A. Scatterplot depicting the relationship of promoter DNA methylation between primary and recurring tumors. Promoters that were differentially methylated between primary and recurring tumors in at least 5 patients are highlighted (DNA methylation difference greater than 75%, adjusted p-value below 0.001, and average RRBS read coverage greater than 20 reads). r: Pearson correlation.
B. Barplots (top) depicting the number of patients that show significant gain or loss of DNA methylation in the differentially methylated promoters highlighted in panel A; scatterplots and line plots (bottom) showing the change in DNA methylation associated with disease progression (measured as percentage points, pp) for patients following the cohort-trend (red) or not (blue). Trend lines were calculated using the *loess* method.
C. Definition of “trend” and “anti-trend” patients based on the Manhattan distance between the maximal trend at differentially methylated promoters (DNA methylation values of 0% or 100%) and the observed difference in DNA methylation for each patient. “Trend” patients are those whose DNA methylation profiles are similar to the maximal trend (low normalized Manhattan distance); “Anti-trend” patients are those whose methylation profiles are most different from the maximal trend (high normalized Manhattan distance).
D. Pathway enrichment analysis of those genes that recurrently lose DNA methylation during disease progression and those that recurrently gain DNA methylation during disease progression.
E. Kaplan-Meier plots displaying progression-free survival and overall survival probabilities over time for patients stratified by whether they followed the cohort trend of differential promoter DNA methylation (trend patients) or not (anti-trend patients), according to the definition in panel C.
F. Kaplan-Meier plots displaying progression-free survival and overall survival probabilities over time for patients stratified into the top-30% patients with increasing or decreasing average DNA methylation levels at the promoters of Wnt signaling genes during disease progression.

Pathway analysis identified an enrichment of genes involved in development and apoptosis among those genes whose promoters gained DNA methylation during disease progression; in contrast, genes whose promoters lost DNA methylation were enriched in the Wnt signaling pathway and T cell activation (Figure 6d). Corroborating the latter finding, when we classified all patients according to whether they on average gained or lost DNA methylation in the promoters of Wnt signaling genes, we observed a significant association between loss of DNA methylation and reduced PFS and OS (Figure 6f).

## Discussion

Focusing on glioblastoma as one of the genetically most complex cancers, we sought to determine the prevalence and character of epigenetic tumor heterogeneity in time and space. To that end, we established a comprehensive set of DNA methylation profiles covering primary and recurring tumors from the same patients as well as multi-sector samples in a subset of patients. A longitudinal cohort of 112 patients with *IDH*-wildtype primary glioblastoma that had undergone at least two (and up to four) tumor resections was assembled based on the Austrian Brain Tumor Registry, thus providing a population-scale representation of glioblastoma patients. An optimized RRBS protocol allowed us to work with minute amounts of FFPE tissue, while providing single-CpG and single-allele resolution and insights into epigenetic heterogeneity at single-cell level that would be difficult or impossible to obtain using microarray-based methods for DNA methylation profiling.

Based on the RRBS dataset, we were able to infer a broad range of tumor properties – including glioblastoma transcriptional subtypes, aspects of the tumor microenvironment, and tumor cell-intrinsic attributes such as cell proliferation. Our analysis revealed changes in tumor microenvironment between primary and recurring tumors and identified the composition of the tumor microenvironment to be a major discriminatory factor between transcriptional subtypes as well as MR progression types, suggesting potential clinical applications of DNA methylation based prediction of the tumor microenvironment. In line with recent work^13,20^ we observed co-occurrence of multiple transcriptional subtypes within the same tumor and frequent switching of the dominant subtype over time. Moreover, patients whose tumor switched to the mesenchymal subtype had reduced survival. Assessing the regulatory basis of the transcriptional subtypes, we identified epigenomic signatures of increased EZH2 activity in glioblastoma of the mesenchymal subtype. In light of the significantly worse prognosis of mesenchymal tumors, these results might enable new subtype-specific therapeutic approaches that exploit the observed regulatory differences, such as the use of emerging EZH2 inhibitors^49-51^.

We also observed characteristic trends in DNA methylation between primary and recurring tumors, including a demethylation of Wnt signaling gene promoters that was associated with worse prognosis. Aberrant activation of the Wnt signaling pathway is observed in various cancers including glioblastoma, where hypermethylation mediated suppression of Wnt signaling inhibitors was identified as a source of aberrant activation of this pathway^52^. We quantified epigenetic tumor heterogeneity in two complementary ways, and DNA methylation erosion as measured by the PDR score was associated with survival. Patients whose primary tumors harbored higher levels of DNA methylation erosion showed longer PFS and a tendency towards longer OS. These results were surprising given that previous studies in hematopoietic malignancies (AML^26^ and CLL^27^) had associated increased epigenomic heterogeneity with worse prognosis. This discrepancy might be explained by the fact that chemotherapy in leukemia is applied to the entire population of malignant cells, while the bulk of glioblastoma is surgically removed prior to chemotherapy and radiotherapy. The non-selective bottleneck of tumor resection might turn the evolutionary advantage of a heterogeneous population of tumor cells into a disadvantage, especially in the case of stochastic and therefore mostly detrimental DNA methylation erosion.

Finally, we observed that several properties of the recurring tumors were specifically associated with survival, while there was no strong association in the primary tumors. This was true for transcriptional subtypes (Figure 2e), MR-imaging derived necrotic and enhancing tumor volumes (Supplementary Fig. 4e), CD68+ macrophage infiltration (Figure 3c), and tumor cell proliferation levels (Figure 4b). These results emphasize the potential clinical relevance of repeated biopsy and detailed diagnostic work-up of recurring tumors in order to promote more personalized treatment decisions upon glioblastoma recurrence.

In summary, our study establishes a rich resource describing the DNA methylation dynamics of glioblastoma progression in a highly annotated clinical cohort with matched MR imaging and detailed histopathological analyses that included the tumor microenvironment. Importantly, all data are openly available through public repositories and a detailed Supplementary Website (http://glioblastoma-progression.computational-epigenetics.org/). This study also highlights the feasibility and potential of working with national patient registries and large patient cohorts, with FFPE samples, and with clinical data that have been collected as part of routine clinical care. Finally, in combination with research that established the accuracy and robustness of DNA methylation assays for clinical diagnostics^54^, our data support that DNA methylation sequencing can make a relevant contribution to the clinical assessment of tumor heterogeneity, providing potential biomarkers for improved diagnosis, prognosis, and personalized therapy in glioblastoma and other heterogeneous cancers.

## Methods

### Sample acquisition via a population-based registry

All glioblastoma cases were selected from the Austrian Brain Tumor Registry^34^, including only patients over 18 years of age with a first surgery at diagnosis and at least one additional surgery upon recurrence. Tumor samples and clinical data were provided by the following partner institutions: Medical University of Vienna, Kepler University Hospital Linz, Paracelsus Medical University Salzburg, Medical University of Innsbruck, Karl-Landsteiner University Hospital St. Pölten, State Hospital Klagenfurt, State Hospital Wiener Neustadt, Hospital Rudolfstiftung Vienna, and the Medical University of Graz. The resulting cohort comprised 159 patients with matched FFPE samples for the primary tumor and at least one recurring tumor, which were deposited in the neurobiobank of the Medical University of Vienna (ethics vote EK078-2004). After screening for sufficient tumor content in both the primary and the recurring tumor, 47 patients were excluded. A total of 112 patients were retained, each with at least two and up to four time points (283 tumor samples in total, including 6 patients with multi-sector sampling). The diagnosis of primary glioblastoma, *IDH*-wildtype was confirmed by central pathology review according to the 2016 update of the WHO classification^55^ including targeted assessment of the IDH R132H mutational status. In addition, 13 patients (32 tumor samples) with IDH-mutant oligodendroglioma/astrocytoma/glioblastoma, and 5 patients (5 samples) who underwent temporal lobe surgery due to epilepsy (Medical University of Vienna) were included as controls. Informed consent was obtained according to the Declaration of Helsinki, and the study was approved and overseen by the ethics committee of the Medical University of Vienna (ethics votes EK550/2005, EK1412/2014, EK 27-147/2015).

### DNA isolation from FFPE tumor samples

Areas of highest tumor cell content were selected based on hematoxylin-eosin stained sections, while any samples with tumor cell content below 50% in the region-of-interest were excluded from further analysis. Genomic DNA was extracted from FFPE tissues using the QIAamp DNA FFPE Tissue Kit following manufacturer’s instructions.

### DNA methylation profiling by RRBS

RRBS was performed as described previously^29^ using 100 ng of genomic DNA for most samples, while occasionally going down to 2 ng (if not more DNA was available) and up to 200 ng (Supplementary Table 2). To assess bisulfite conversion efficiency independent of CpG context, methylated and unmethylated spike-in controls were added in a concentration of 0.1%. DNA was digested using the restriction enzymes MspI and TaqI in combination (as opposed to only MspI in the original protocol) in order to increase genome-wide coverage. Restriction enzyme digestion was followed by fragment end repair, A-tailing, and adapter ligation. The amount of effective library was determined by qPCR, and samples were multiplexed in pools of 10 with similar qPCR *C_t_* values. The pools were then subjected to bisulfite conversion followed by library enrichment by PCR. Enrichment cycles were determined using qPCR and ranged from 12 to 21 (median: 16). After confirming adequate fragment size distributions on Bioanalyzer High Sensitivity DNA chips (Agilent), libraries were sequenced on Illumina HiSeq 3000/4000 machines in a 50 or 60 basepair single-read setup.

### DNA methylation data processing

RRBS data were processed using a custom pipeline based on Pypiper (http://databio.org/pypiper) and Looper (http://databio.org/looper). Adapter sequences were trimmed, and 60 basepair reads were cropped to 50 base-pairs using Trimmomatic^56^ with the following settings: ILLUMINACLIP:RRBS_adapters.fa:2:40:7 SLID-INGWINDOW:4:15 MAXINFO:20:0.50 CROP:50 MINLEN:18. Trimmed reads were then aligned to the human genome build hg38 using BSMAP in RRBS mode^57^, and DNA methylation calling was performed with a custom python script (biseqMethCalling.py) published previously^29^. To assess bisulfite conversion efficiency, unmapped reads were aligned to the spike-in reference sequences using Bismark^58^, and DNA methylation calls for methylated and unmethylated controls were extracted from the alignment file. CpGs in repetitive regions according to the UCSC RepeatMasker track were excluded from further analysis. DNA methylation data were analyzed at the level of single CpGs or in a binned format with mean DNA methylation values calculated across 5-kilobase regions, CpG islands (as defined in the UCSC Genome Browser) or GENCODE promoter regions (1 kilobase upstream to 500 bases downstream of the transcription start site).

### Identification of copy number aberrations from RRBS data

Copy number aberrations for each sample were identified using the R/Bioconductor package CopywriteR^40^ based on the BSMAP-aligned BAM files and a bin size of 100,000. Data from five normal brain controls were merged at the level of aligned bam files to serve as the shared control for all analyses. Each individual sample was then normalized either against the merged control or against the cohort median, whichever showed the less extreme (i.e., more conservative) value for a given bin. Genomic segments identified by CopywriteR were classified as significantly amplified or deleted if their normalized absolute copy number value deviated more than one cohort standard deviation (mean standard deviation across all bins in a given segment) from 0. Significantly amplified or deleted segments for each sample were then plotted in an overview graph sorted by segment length.

### Identification of single nucleotide variants from RRBS data

Single nucleotide variants and small insertions and deletions were identified using Bis-SNP^42^ based on the BSMAP-aligned BAM files, the human reference genome build hg38, and dbSNP build 147. Identified variants were annotated using SnpEff v4.2.

### Annotation of glioblastoma associated genes

Glioblastoma associated genes were taken from a recent publication^59^ and annotated for their cancer-linked function (oncogene, tumor suppressor gene, drug resistance gene) based on a published classification of cancer genes^60^. Genes not contained in this classification were manually annotated according to their known or suspected molecular functions as described in GeneCards (http://www.genecards.org).

### Patient survival analysis

Survival analysis was performed using the functions survfit() and survdiff() of the R package ‘survival’. For continuous variables, patients with the 50% highest values were compared to those with the 50% lowest values (unless indicated otherwise). Survival curves were plotted with ggsurvplot() from the R package ‘survminer’.

### Inference of transcriptional subtypes from RRBS data

Glioblastoma transcriptional subtypes^43^ were predicted from DNA methylation data at the level of single CpGs using L2-regularized logistic regression as implemented in the R package ‘LiblinearR’. Classifiers were trained and evaluated on Infinium 27k DNA methylation data for glioblastoma tumors^44^ obtained from the TCGA data portal (https://portal.gdc.cancer.gov/). TCGA data were restricted to IDH wildtype, non-G-CIMP samples.

Furthermore, neural subtype samples were excluded because this subtype of glioblastoma had previously been associated with tumor margin and contamination with non-tumor brain tissue^61^. For each sample in our cohort, a classifier was trained and evaluated on the TCGA data, using those CpGs that were covered also in the sample of which the transcriptional subtype was to be predicted (1,249 CpGs on average). After performance evaluation by 10-fold cross-validation and calculation of the cross-validated receiver operating characteristic (ROC) area under curve (AUC) values, a final classifier was built using all selected TCGA samples. This classifier was used to predict the transcriptional subtype including class probabilities of the respective sample.

### Groupwise differential DNA methylation analysis

Differentially methylated CpGs between predefined groups of tumor samples were identified with a custom R script that uses a two-sided Wilcoxon rank-sum test. Groups containing less than five samples were excluded from the analysis, and only CpGs covered by at least five reads per sample in at least 30% of samples were included. CpGs in repetitive regions (“RepeatMasker”, “Simple Repeats”, and “WM + SDust” tracks from the UCSC Genome Browser, downloaded 6 September 2016) were also excluded. For the retained CpGs, differential DNA methylation between groups of samples was assessed using the Wilcoxon rank-sum test (wilcox.test() in R), and p-values were adjusted for multiple testing using the Benjamini-Hochberg method (p.adjust() in R). CpGs with multiple-testing adjusted p-values smaller than 0.05 and with a median difference of beta values larger than 0.1 were considered significant.

### Region set enrichment analysis using LOLA

Enrichment of genomic region sets among the differentially methylated regions was assessed using the LOLA software^46^. To reduce potential biases from co-located CpGs, CpGs were merged into 1-kilobase tiling regions across the genome prior to LOLA analysis. In LOLA, the hypermethylated or hypomethylated regions were used as the query set, and the set of all differentially methylated tiling regions were used as the universe. Only regions from astrocytes or embryonic stem cells in the LOLA Core database were included in the analysis for better interpretability. P-values were corrected for multiple testing using the Benjamini and Yekutieli method (p.adjust() in R), and all enrichments with an adjusted p-value below 0.05 were considered significant. In a control experiment, to assess potential effects of imbalance between hypermethylated and hypomethylated region sets, the analysis was repeated using the top-N highest ranking regions from both sets.

### DNA methylation inferred regulatory activity (MIRA)

MIRA scores for selected sets of transcription factor binding sites from the LOLA Core database^46^ were calculated as in the original publication^28^. Briefly, aggregated DNA methylation profiles around the binding sites (2.5 kilobases upstream and downstream, split into 21 bins) were created for each sample and transcription factor. MIRA scores were calculated as the log ratio between aggregated DNA methylation values for the center bin (bin 0, reflecting the binding site) and the average of two flanking bins (bins -5 and +5).

### Immunohistochemistry

The following antibodies were used for immunohistochemistry: IDH1 (1:60 Dianova #DIA-H09), CD3 (1:200 Thermo Scientific #RM-9107-S1), CD8 (1:100 Dako Cytomation #M7103), CD45Ro (1:500 Dako Cytomation #M0742), CD8 (1:100 Dako Cytomation #C8/144B), FoxP3 (1:25 BioLegend #320116), CD163 (1:1000 Novocastra #NCL-L-CD163), CD68 (1:5000 Dako Cytomation #M0814), HLA-DR (1:400 Dako Cytomation #M0775), MIB1 (1:200 Dako Cytomation #M7240), CD34 (1:100 Novocastra #NCL-l-END).

FFPE blocks were cut at a thickness of 3 μm, and sections were stained on a Dako autostainer system using the following primary antibodies MIB1, HLA-DR, CD34, CD45Ro, CD68, CD3, CD8, CD163. Antigens were retrieved by heating the sections in 10 mM sodium citrate (pH 6.0) at 95°C for 20 min, followed by incubation with primary antibodies for 30 min at room temperature. The Dako Flex+mouse detection system was used according to manufacturer’s recommendations. For primary antibodies against IDH1 and FoxP3, a Ventana BenchMark automated staining system was used, followed by visualization using the Ultra View detection kit. All sections were counterstained with hematoxylin. For each antibody used positive and negative controls were used per 30-slide-batch. Negative controls were performed by omitting the primary antibody and by using Universal Negative Control rabbit (Dako) for polyclonal rabbit antibodies or purified mouse myeloma IgG1 (Zymed Laboratories, San Francisco, CA) for monoclonal mouse antibodies. The slides were scanned using a Hamamatsu NanoZoomer 2.0 HT slide scanner, and images were analyzed using the NDPview 2 software. Whole slide scans were downsampled to 5x magnification and exported as JPEG images. Fiji was used for further image processing^62^. First, the inbuilt Color Deconvolution method was used to separate the hematoxylin from the DAB stain. The 8-bit greyscale hematoxylin image was thresholded using Phansalkar thresholding, and nuclei were counted using the built-in Analyze Particles algorithm to determine reference cell counts for each image. For each antibody stain Phansalkar thresholding parameters were manually optimized, followed by automated counting of DAB-positive cells. The inferred counts for antigen-expressing cells and corresponding total nuclei counts were saved as spreadsheet for further statistical analysis.

### Histopathological analysis of whole slide scans

H&E stained slides were scanned using a Hamamatsu NanoZoomer 2.0 HT slide scanner. The Hamamatsu NDP.view2 software was used to annotate relevant regions in the slide scans. Areas including hemorrhages, necrosis, scars, squeezed tissue, and preexisting brain parenchyma were manually segmented by a specialist in neuropathology using the Freehand Region tool. The slide scans were downsampled to 10x magnification and exported as jpeg images using the NDPITools plugin of Fiji^62-64^ along with xml files featuring the annotations (^*^.ndpa). The JPEG images were loaded into Fiji for further processing.

To obtain a cell nuclei mask for each slide scan image, the Color Deconvolution method of Fiji was used to obtain an 8-bit greyscale image of the hematoxylin stain^65^. Automated local thresholding based on Phansalkar’s method segmented cell nuclei (parameters k = 0.2, r = 0.5, radius = 8)^66^, followed by the binary Close and Open operations. The Watershed method was used to separate clustered nuclei^67^. To obtain a mask of the gross tissue on the slide scan, the original image was first converted to an 8-bit greyscale image followed by global thresholding (thresholding values 0-207). The masks were loaded into MATLAB R2014b (MathWorks) for further automated image analysis. For each image, the annotation data was fetched from the respective xml file and converted into polygons. These polygons were subsequently grouped to form binary masks based on their annotation (e.g., a necrosis mask comprising all annotated necrotic areas). The gross tissue was determined by first loading the previously obtained gross tissue mask into MATLAB, performing the binary Close operation (dilating and eroding the binary image by 30 pixels) and then removing all connected components smaller than 1,000 pixels. Pixels that were neither included in the background nor in any of the image annotation masks were assigned to the tumor. Using the pixel-to-area conversion, the areas covered by tumor, necrosis and other annotated tissues were calculated.

Subsequently, the pre-obtained binary nucleus mask was analyzed in blocks of 160 × 160 pixels (~146 μm × 146 μm). This block size was empirically found to provide a good compromise between spatial resolution and nuclear content required for statistical analysis. The centroids of the nuclei were localized and the area and eccentricity (which is calculated as the distance between the foci of a fitted ellipse divided by its major axis length, with values close to 0 corresponding to circular nuclei and values close to 1 corresponding to spindle-shaped nuclei) of each localized nucleus was assessed. Next, the number of nuclei per block was calculated from the centroid map. From the nucleus areas in each block, the average area and eccentricity as well as the standard deviation of the nucleus areas and eccentricities were determined. The coefficient of variation was calculated by dividing the standard deviation by the average for each block. For nuclei number, average area and eccentricity, standard deviation and coefficient of variation, the data from all blocks were assembled in a matrix and saved as grey scale bitmap (.bmp) image as well as color portable network graphics (.png) image. Furthermore, using the binary annotation masks, each block was assigned to its corresponding region, given that >90% of the pixels in that block were uniformly annotated. Subsequently, for each type of region, the mean, standard deviation, coefficient of variation, median, and mode of the aforementioned nucleus characteristics were calculated (e.g., obtaining the mean average nucleus density in tumor tissue). For further statistical processing, the numerical data extracted from the slide scan images were saved into a spreadsheet.

### Radiological evaluation of glioblastoma patients

MR images of glioblastomas at time of first diagnosis and recurrence in sufficient quality were available for 54 of the glioblastoma patients included in this study, which were contributed by 6 different radiology departments. Both T1-weighted images with contrast enhancement (CE) and fluid-attenuated inversion recovery (FLAIR)/T2-weighted axial images were reviewed for topographic tumor location to assess solitary versus multicentric tumors and local versus distant recurrences. Multicentric glioblastomas were defined as at least two spatially distinct lesions that are not contiguous with each other and whose surrounding abnormal FLAIR/T2 signals do not overlap^68^. Tumor segmentation was performed with BraTumIA^69,70^, which uses multi-modal MRI sequences for fully automated volumetric tumor segmentation. T1, T1 contrast enhanced, T2, and FLAIR sequences were used to segment four tumor tissue types: necrotic, cystic, edema/non-enhancing, and enhancing tumor. Due to differences in MRI protocols across the study sites, the multi-modal sequences were affine registered to the T1 sequence with SPM122 and resampled to 1x1x3mm voxel size prior to segmentation. The BraTumIA-derived segmentations were reviewed by an expert radiologist, and errors in the automatic segmentation were manually corrected.

### Evaluation of MR imaging-based progression phenotypes

MR imaging-based tumor progression was assessed according to the Response Assessment in Neuro-Oncology (RANO) standard^71^. Serial T1-weighted images with CE and FLAIR/T2-weighted images were available for 43 patients. Progression subtypes were classified as described previously^33,72^: (i) Classic T1 (incomplete disappearance of T1-CE during therapy followed by T1-CE increase at progression), (ii) cT1 relapse / flare-up (complete disappearance of T1-CE during therapy followed by T1-CE reoccurrence at progression), (iii) Primary non-responder (increase and/or additional T1-CE lesions at first MR imaging follow-up after start of therapy), (iv) T2-circumscribed (bulky and inhomogeneous T2/FLAIR progression, no or single faintly speckled T1-CE lesions at progression), and (v) T2-diffuse (complete decrease in T1-CE during therapy but exclusive homogeneous T2/FLAIR signal increase with mass effect at progression).

### DNA methylation based prediction of tumor properties

Tumor properties such as the immune cell infiltration and tumor cell morphology were predicted from DNA methylation data using a machine learning approach that was based on the R package ‘LiblinearR’. The DNA methylation data were prepared by calculating for each sample the mean DNA methylation levels in 5-kilobase tiling regions across the genome. Tiling regions covered in less than 90% of the samples were excluded from the analysis, and the filtered data matrix (samples x tiling regions) was subjected to imputation using the function impute.knn() from the R-package ‘impute’, with the parameter k (i.e., the number of nearest neighbors considered) set to 5. Tumor properties represented by continuous response variables were converted into categorical variables by setting the 20% highest values to ‘high’, the 20% lowest values to ‘low’, and the remaining samples to ‘NA’. Imputed beta values were used to train and evaluate the classifiers using LiblineaR(). In the confirmatory hierarchical clustering based on the most predictive features identified by the classifiers, the beta values were scaled across samples for better visualization and comparability. LiblineaR() was set to use support vector classification by Crammer and Singer as model type, and the appropriate cost parameter was estimated from the imputed data matrix using the function heuristicC() from the same package. For each tumor property, the performance of the classifiers was determined through leave-one-out cross-validation, and 10 control runs with randomly shuffled labels were included to detect potential overfitting. ROC curves and ROC AUC values were determined using the functions prediction() and performance() of the R-package ‘ROCR’. Finally, we trained a classifier on the entire dataset using the selected model and cost parameter, which was then used for further analysis including the extraction of the most predictive features, hierarchical clustering, and the prediction of additional samples (for the transcriptional subtypes).

### Estimation of DNA methylation heterogeneity

The epi-allele entropy as a measure of sub-clonality within a tumor was calculated using a slightly modified version of methclone^48^. We calculated epi-allele entropies separately for each of the samples and, independently, for each matched pair of primary and recurring tumors. Input files to methclone were created by aligning the trimmed RRBS reads to the human reference genome build hg38 using Bismark^58^. As in the original publication, methclone was set to require a minimum of 60 reads in order to consider a locus, and loci with a combinatorial entropy change below -80 were classified as epigenetic shift loci (eloci) between primary and recurring tumors^48^. For each pair, we then calculated epi-allele shifts per million loci (EPM), dividing the number of eloci by the total number of assessed loci normalized to 1 million loci^48^.

The proportion of discordant reads (PDR) as a measure of local erosion and DNA methylation disorder was calculated as described in the original publication^27^. Briefly, the number of concordantly or discordantly methylated reads with at least four valid CpG measurements was determined for each CpG using a custom python script. The PDR at each CpG was then calculated as the ratio of discordant reads compared to all valid reads covering that locus. CpGs at the end of a read were disregarded to remove potential biases due to the end-repair step of RRBS library preparation. Because the PDR and epi-allele entropy calculation is highly sensitive to differences in the read composition of the underlying RRBS library, we focused this analysis on RRBS libraries with a similar number of enrichment cycles (13-15) to ensure high consistency between samples (Supplementary Fig. 6a).

Sample-wise PDR and epi-allele entropy values were calculated by averaging across promoters that were covered in more than 75% of the samples. Promoter regions were defined as the genomic region 1 kilobase upstream to 500 basepairs downstream of a given transcription start site as annotated by GENCODE^73^.

### Pairwise differential DNA methylation analysis

Differentially methylated CpGs between sample pairs (i.e., primary tumor versus matched recurring tumor) were identified with a custom R script that uses Fisher’s exact test. This test was applied to the methylated and unmethylated read counts derived from the BSMAP-aligned reads by the biseqMethCalling.py script. P-values were adjusted for multiple testing using the Benjamini-Hochberg method. To obtain promoter-wise differential DNA methylation calls, p-values were combined using a generalization of Fischer’s method^74^ as implemented in RnBeads^75^. Promoter methylation levels for each sample were calculated as the mean of all CpGs in the promoter region. Promoter regions were defined as the genomic region 1 kilobase upstream to 500 basepairs downstream of a given transcription start site as annotated by GENCODE^73^.

### Pathway enrichment analysis

Enrichment analysis for gene sets and pathways was performed using enrichR^76,77^ through an R interface (https://github.com/definitelysean/enrichR) querying the Panther_2016 database (http://www.pantherdb.org/) for enrichments with an adjusted p-value below 0.05.

### Data and code availability

All data are available through the Supplementary Website (http://glioblastoma-progression.computational-epigenetics.org/). Genome browser tracks facilitate the locus-specific inspection of the DNA methylation data, and a d3.js based graphical data explorer enables interactive analysis of associations in the annotated dataset (Supplementary Fig. 7). The Supplementary Website also hosts the raw and segmented image data from the histopathological analysis as well as the raw and segmented MR imaging data. The processed DNA methylation data will also be available for download from NCBI GEO (accession number: GSE100351, reviewer link: https://www.ncbi.nlm.nih.gov/geo/query/acc.cgi?acc=GSE100351,login token: ufahcwkajrqnhef), and the raw sequencing data will be available from EBI EGA. Finally, in the spirit of reproducible research^78^ the Supplementary Website makes the source code underlying the presented analyses publicly available.

## Acknowledgements

We thank all patients who have donated their samples for this study. We also thank Gloria Wilk, Martina Muck, Susanne Schmid. and Ulrike Andel for technical assistance with immunohistochemical stainings, macrodissection and tumor tissue shavings, Simon Mages for contributing to the interactive data visualization, the Biomedical Sequencing Facility at CeMM for assistance with next generation sequencing, and all members of the Bock lab for their help and advice.

The study was funded in part by an Austrian Science Fund grant (FWF KLI394) to AW, a Marie Curie Career Integration Grant (European Union’s Seventh Framework Programme grant agreement no. PCIG12-GA-2012-333595) to CB, and an ERA-NET project (EpiMark FWF I 1575-B19). C.B. is supported by a New Frontiers Group award of the Austrian Academy of Sciences and by an ERC Starting Grant (European Union’s Horizon 2020 research and innovation programme, grant agreement no. 679146). ABTR activities are further supported by unrestricted research grants of Roche Austria to JAH and the Austrian Society of Neurology to SO.

## Author contributions

JKl, AW, and CB designed the study. BK, TR, NP, KHN, JF, MN, MA, MM, TS, GL, BB, JAH, and AW established and annotated the clinical cohort. AK and PD performed DNA methylation profiling. JKl performed the data analysis with contributions from NF and NCS, PM, CFF, JKe, AEG, GS, MK, SO, FM, SW, JT, JB, JPi, JH, SK, KMA, GvC, FP, CS, JPr, PAW, WK, FW, TBK, MS, SS, KD, MP, EK, GW, and CM contributed tumor samples and clinical data. JKl, AW, and CB wrote the manuscript with contributions from all authors.

## Supplementary Tables

**Supplementary Table 1. Patient summary table**

**Supplementary Table 2. RRBS summary table**

### Supplementary Figures

**Supplementary Figure 1.**
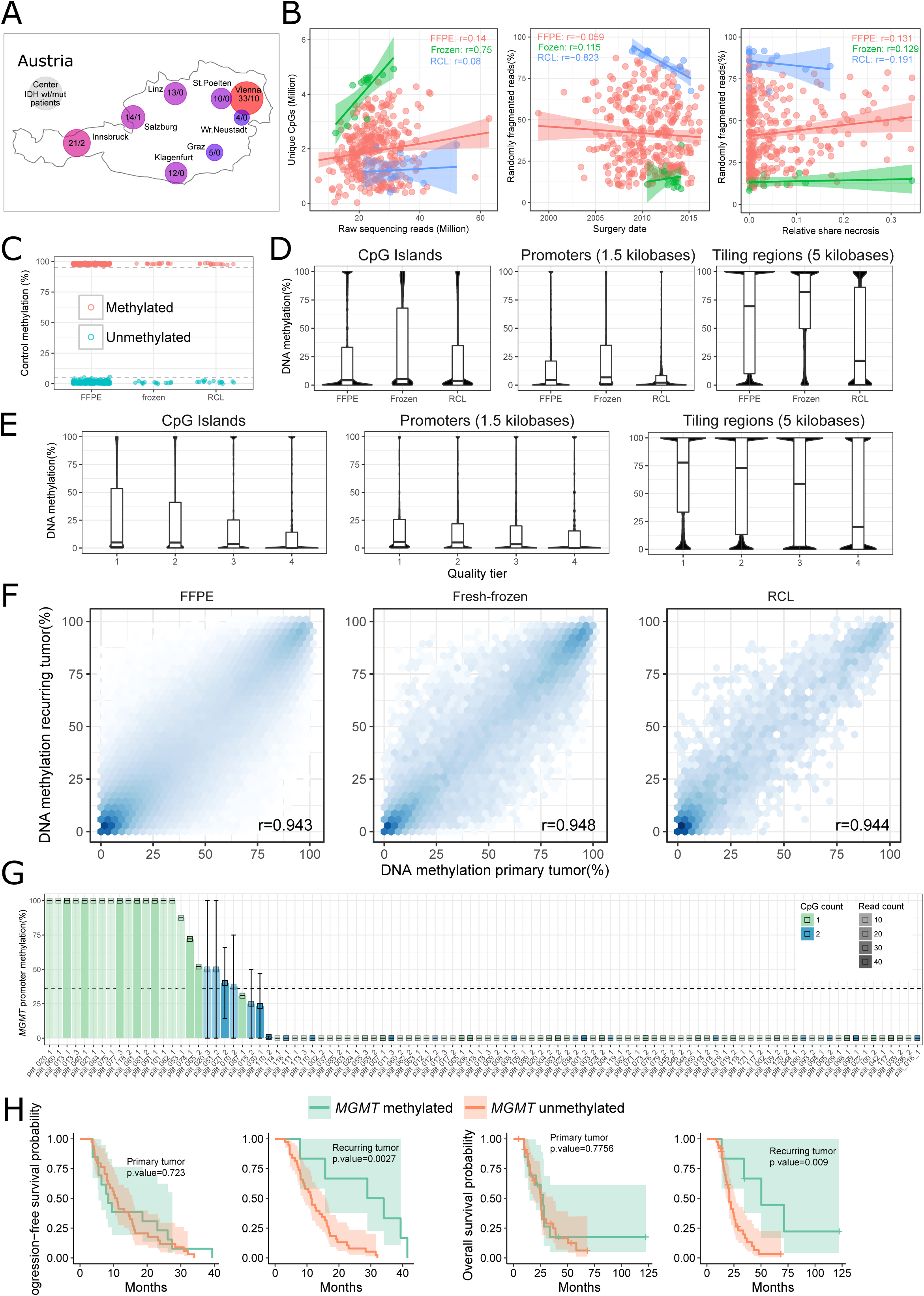
RRBS profiling in a population-based glioblastoma cohort. A Overview of the clinical centers that contributed to this study. The numbers of IDH wildtype and IDH mutated patients are indicated for each center.
B Scatterplots summarizing the RRBS sequencing data. The proportion of randomly fragmented reads (i.e., reads not starting with the expected RRBS restriction sites) reflect the degree of pre-fragmentation of the input DNA. The different sample types (FFPE: formalin-fixed paraffin-embedded; Frozen: fresh-frozen, RCL: ethanol-based conservation) are indicated by color.
C DNA methylation levels of methylated and unmethylated synthetic spike-in control sequences. Dashed lines indicate DNA methylation levels of 5% and 95%.
D, E Distribution of DNA methylation levels across different genomic regions (covered by more than 10 reads per CpG) and for the different sample types (D) and quality tiers (E) defined by the number of unique CpGs detected in each RRBS library (tier 1: more than 3 million; tier 2: between 2 and 3 million; tier 3: between 1 and 2 million; tier 4: below 1 million).
F Scatterplots depicting the relationship of DNA methylation levels in 5-kilobase tiling regions (containing more than 25 CpGs and covered by more than 10 reads per CpG) between primary and recurring tumors for the three different sample types. r: Pearson correlation.
G Mean *MGMT* promotor methylation levels averaged across two CpGs (cg12434587 and cg12981137)^53^. Error bars indicate the maximum and minimum detected methylation levels in each samples. The dashed line indicates the threshold (36%) below which samples are considered unmethylated.
H Kaplan-Meier plots displaying progression-free survival and overall survival probabilities over time for patients stratified by *MGMT* promoter methylation status as depicted in panel G.

**Supplementary Figure 2.**
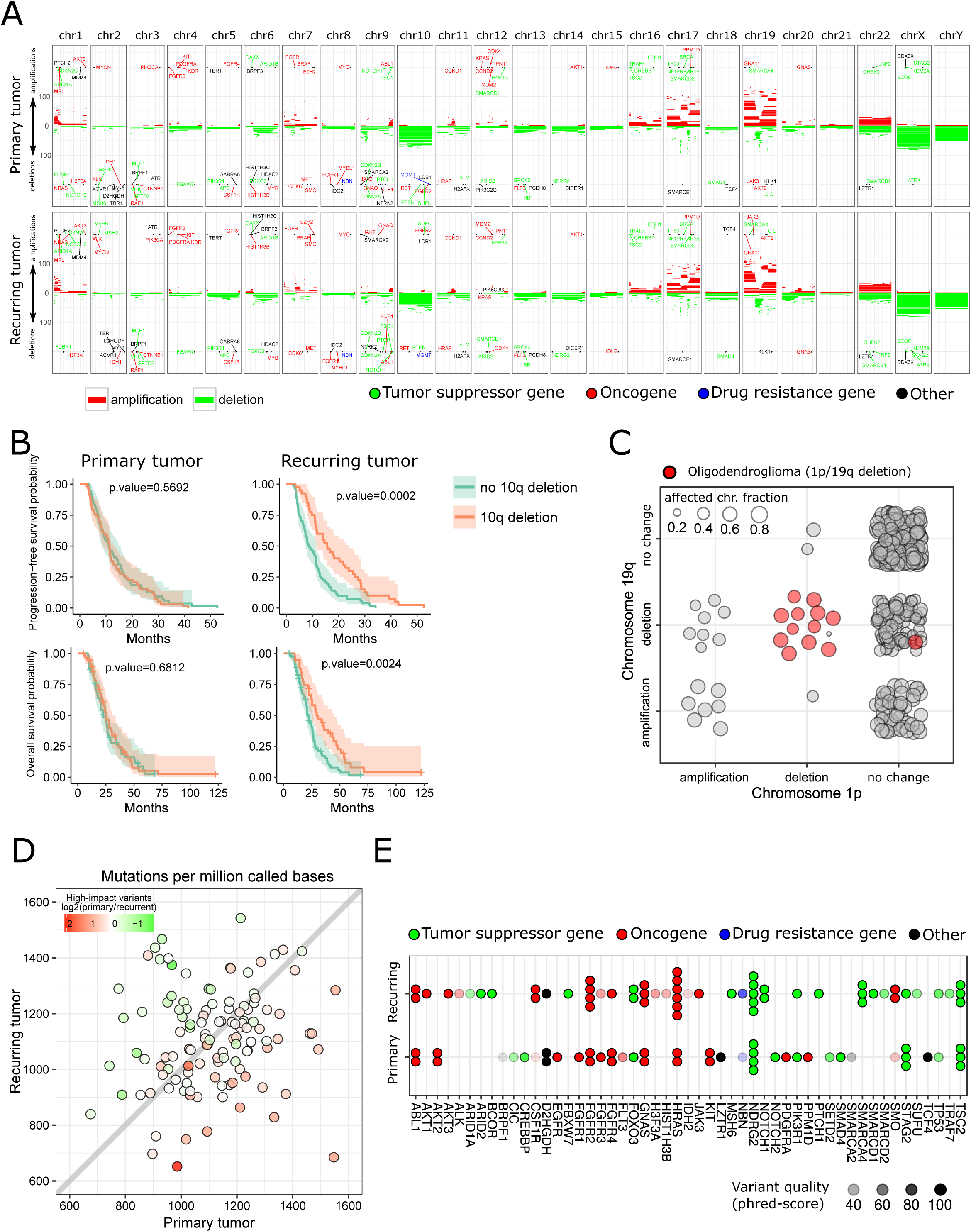
Inference of genetic information from RRBS data. A. Overview of the RRBS-based inference of copy number aberrations (CNAs) in the glioblastoma cohort. The horizontal red and green lines represent the aberrations identified in each of the samples. The genomic position of genes with reported relevance in glioblastoma are indicated.
B. Kaplan-Meier plots displaying progression-free survival and overall survival probabilities over time for patients stratified by chromosome 10q deletion status as depicted in panel A.
C. Assessment of 1p/19q co-deletion status in the IDH wildtype primary glioblastoma samples as well as the 13 oligodendroglioma samples from 6 patients with known 1p/19q co-deletion as positive controls. The size of the bubbles represents the mean fraction of the respective chromosome arms that are affected by the indicated CNAs.
D. Scatterplot displaying the relationship of normalized RRBS-derived mutation calls (SNPs and InDels) between primary and recurring tumors.
E. Cohort-wide mutational profile of genes with known relevance in glioblastoma. Only mutations with high predicted impact on protein function are displayed. Each dot represents a tumor sample in which the indicated mutation was detected. Variant qualities above 100 were set to 100.

**Supplementary Figure 3.**
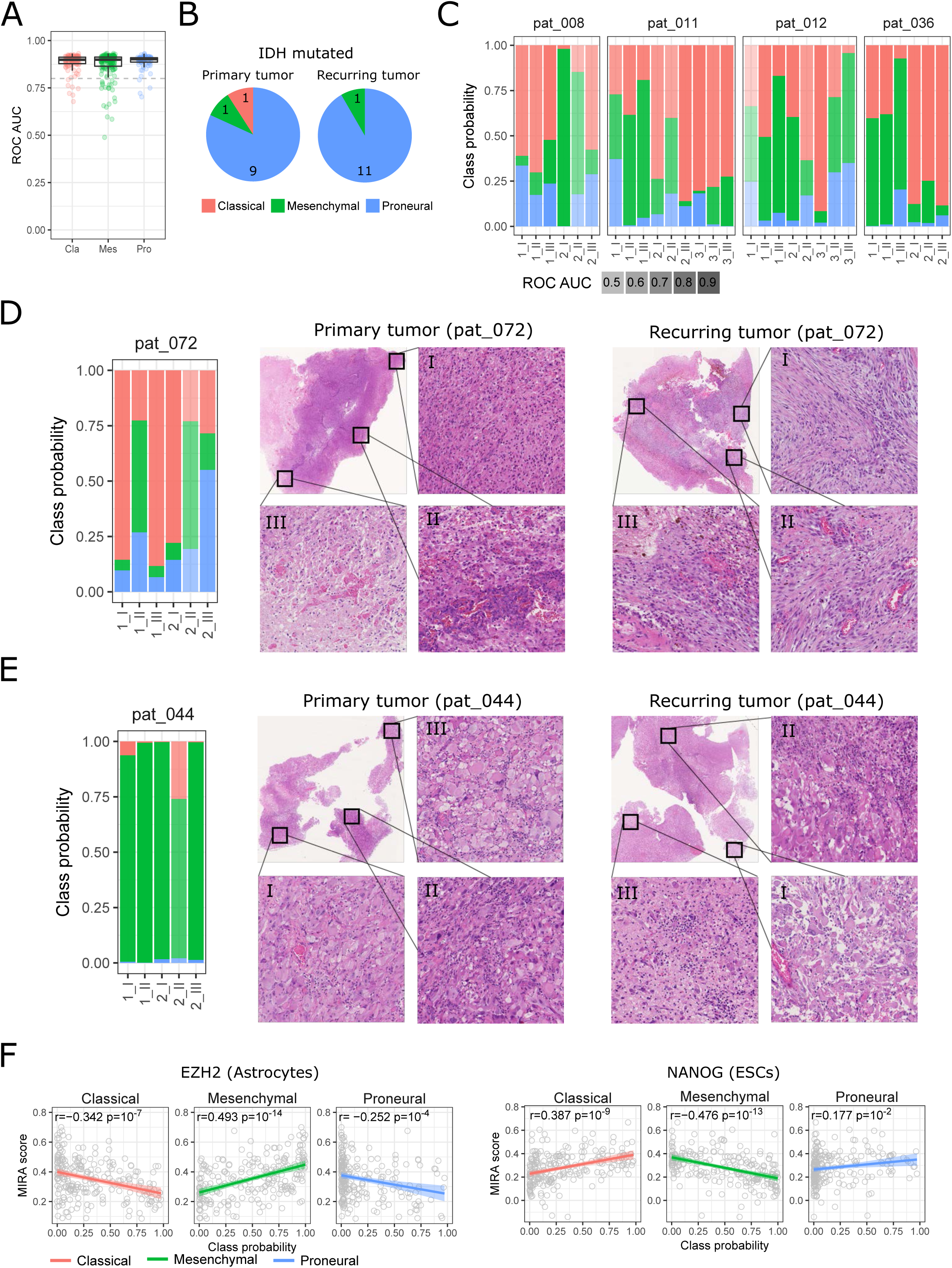
Prediction of glioblastoma transcriptional subtypes from RRBS data. A Sample-wise ROC AUC values as a measure for the accuracy of transcriptional subtype prediction from RRBS data. Dashed line: ROC AUC = 0.8
B Transcriptional subtype distribution for IDH mutated samples. The number of samples assigned to each subtype is indicated.
C, D, E Distribution of class probabilities across different regions of the same tumor (indicated by Roman numbers) and across different surgeries (indicated by Arabic numbers) for four out of six patients with multisector samples (C) and the remaining two patients (D,E) accompanied by matched hematoxylin and eosin stains to display the tumor regions from which the multi-sector samples originated. Section III of the primary tumor of patient 44 did not yield enough CpGs to support confident classification.
F Scatterplots displaying the relationship between MIRA scores for the indicated factors and the class probabilities of the indicated transcriptional subtypes. r: Pearson correlation, p: p-value.

**Supplementary Figure 4.**
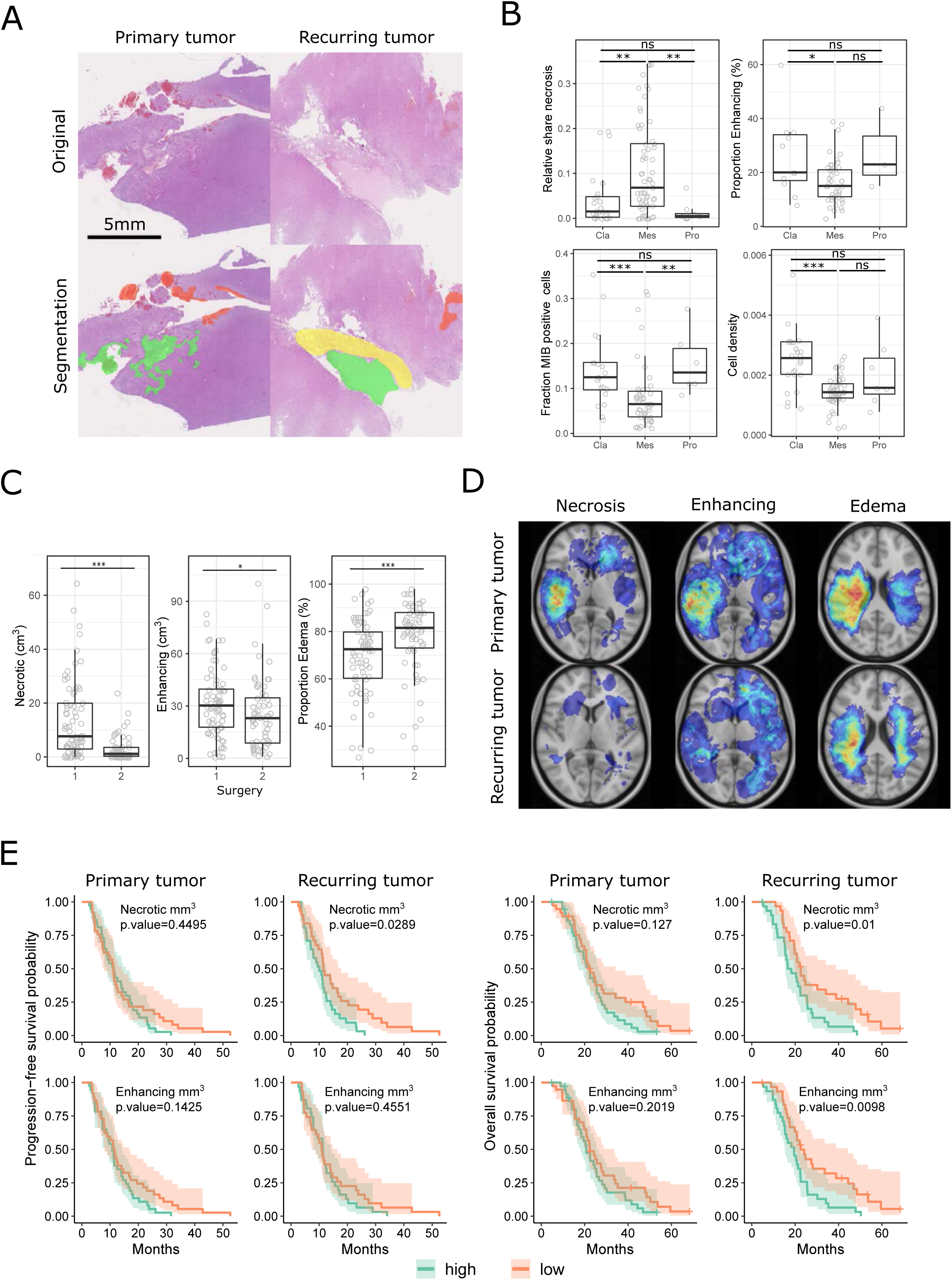
Association of histopathological and MR imaging-derived tumor properties with transcriptional subtypes and disease progression. A. Segmented hematoxylin and eosin stains illustrating the quantification of the histopathological tumor properties (red: hemorrhage, green: necrosis, yellow: meningeal scarring).
B. Comparison of histopathological tumor properties between the different transcriptional subtypes (Cla: classical, Mes: mesenchymal, Pro: proneural).
C. Comparison of histopathological tumor properties between first surgery (primary tumor) and second surgery (recurring tumor).
D. Segmented MR imaging pictures illustrating the quantification of the different MR imaging-derived tumor properties. Heatmap intensity overlays indicate the extent of necrosis, contrast-enhancing (active) tumor volume, and edema in the entire cohort.
E. Kaplan-Meier plots displaying progression-free survival and overall survival probabilities over time for patients stratified according to the level of necrotic and contrast-enhancing (active) tumor volume as derived from MR imaging in primary and recurring tumors. All significance tests comparing groups of samples in this figure were performed using a two-sided Wilcoxon rank sum test: ^*^ p-value < 0.05, ^**^ p-value < 0.01, ^***^ p-value < 0.001, ns: not significant.

**Supplementary Figure 5.**
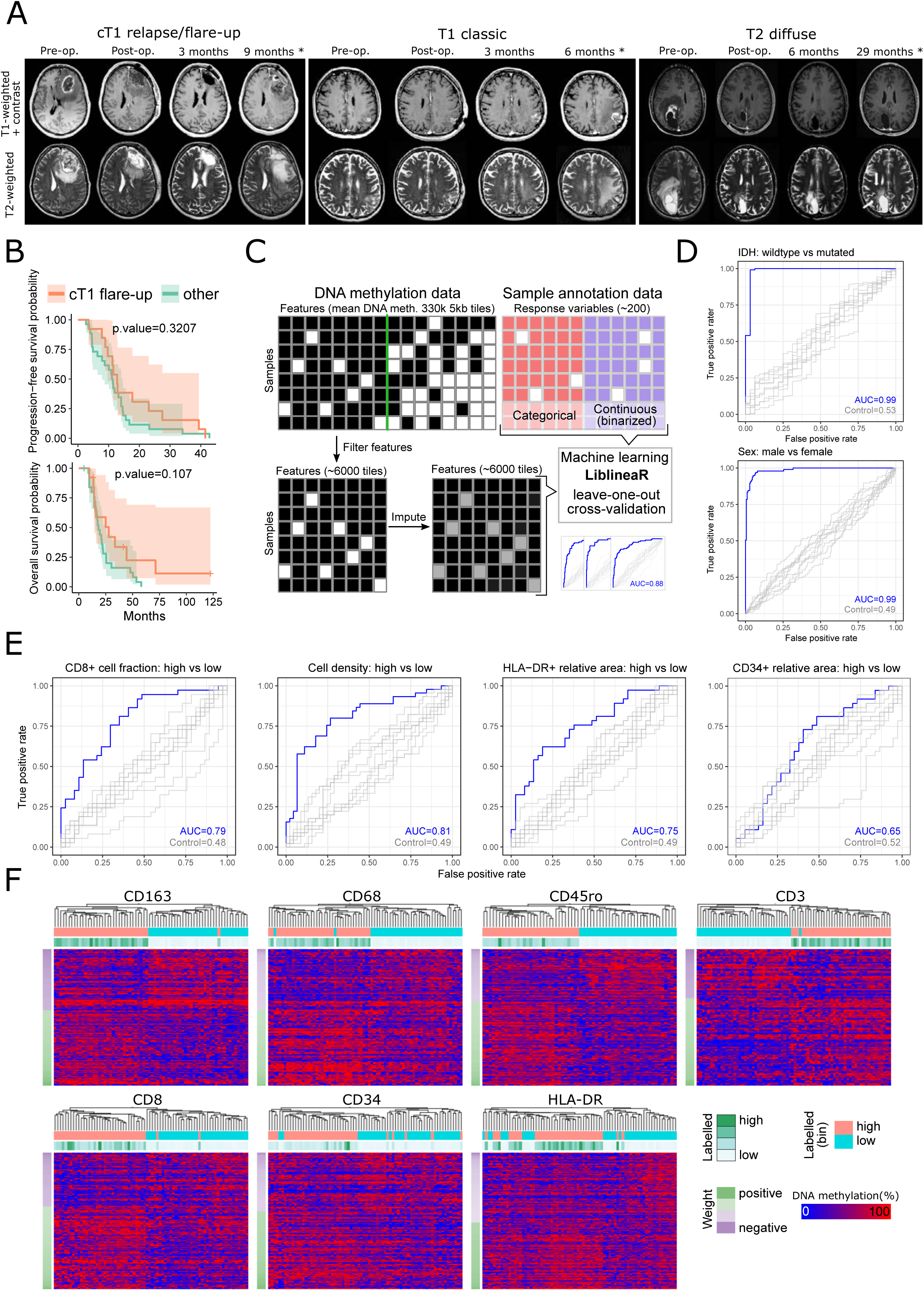
MR imaging progression types and DNA methylation based prediction of tumor properties. A. T1-contrast enhanced and T2/FLAIR MR sequences at each follow-up visit illustrating the three MR imaging progression types in this cohort (cT1 relapse / flare-up, classic T1, T2 diffuse). ‘^*^’: tumor recurrence.
B. Kaplan-Meier plots displaying progression-free survival and overall survival probabilities over time for patients stratified according to their MR imaging progression types.
C. Schematic illustrating the machine learning approach (including the data pre-processing) used to assess the predictability of various tumor properties from RRBS DNA methylation data. White squares indicate missing values; grey squares indicate imputed values.
D. ROC curves showing high prediction accuracy based on DNA methylation data for two features with high expected predictability (IDH mutation status, patient sex).
E. ROC curves evaluating the DNA methylation based prediction of several histopathologic tumor properties.
F. Hierarchical clustering based on the column-scaled DNA methylation values of the most predictive features (5-kilobase tiling regions) as identified by the machine leaning classifiers built to predict the infiltration levels of the indicated immune cell types (cells positive for CD163, CD68, CD45ro, CD3, or CD8) or the extent of indicated tumor properties (cells positive for CD34 or HLA-DR).

**Supplementary Figure 6.**
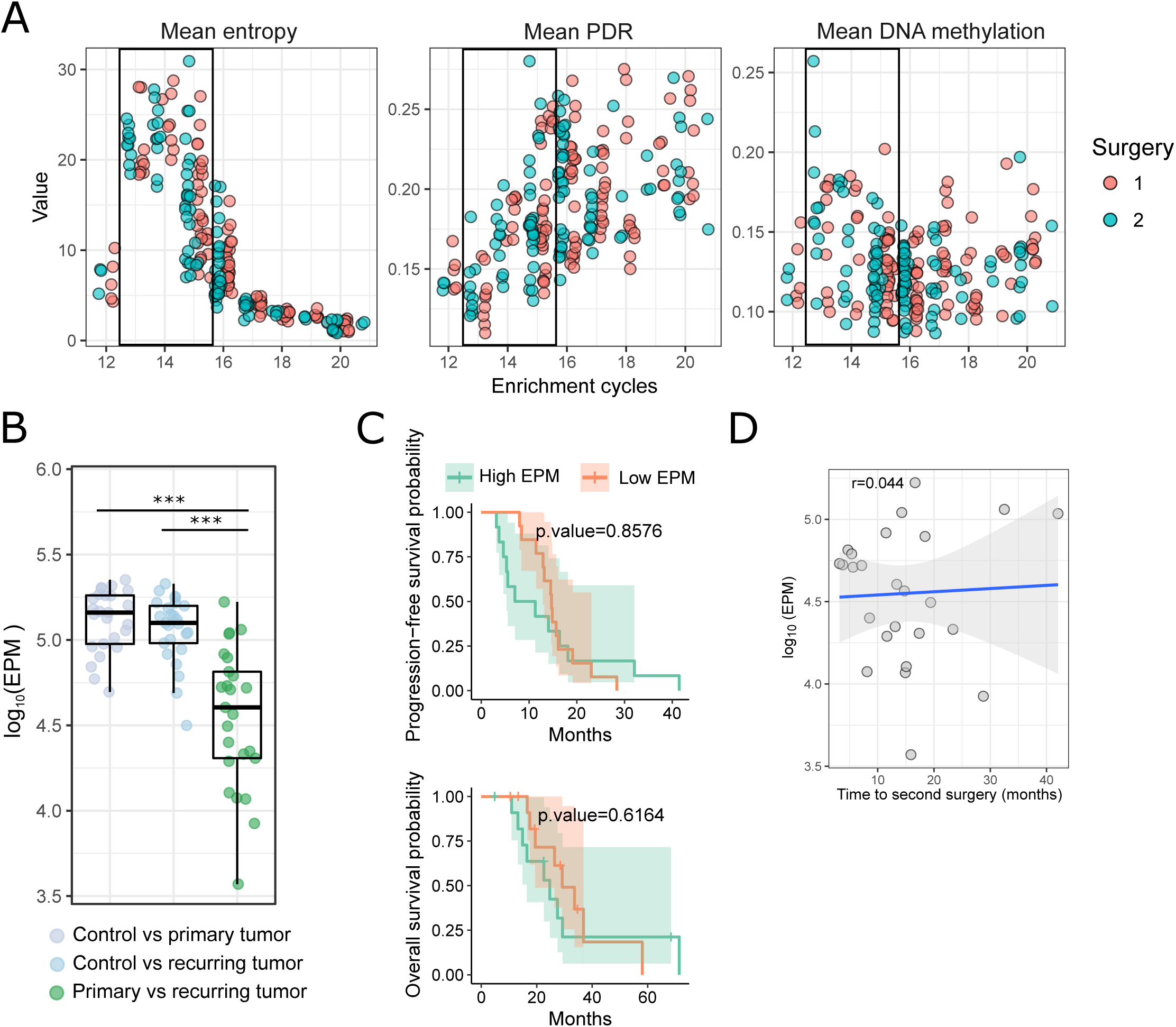
Analysis of DNA methylation heterogeneity in primary and recurring tumors. A. Scatterplots displaying the relationship between PCR enrichment cycles during RRBS library preparation and the indicated measures of DNA methylation heterogeneity. In order to reduce the effect of technical variability, we limited the analysis of epigenomic heterogeneity to samples that fall into a defined narrow range of PCR enrichment cycles (13-15 cycles, indicated by black boxes).
B. Degree of epi-allelic shifting between normal brain control and primary or recurring tumors, as well as between primary and recurring tumors measured by the relative number of loci that show high changes in epi-allele composition (EPM: eloci per million assessed loci). ^***^: p-value < 0.001 (two-sided Wilcoxon rank sum test)
C. Kaplan-Meier plots displaying progression-free survival and overall survival probabilities over time for patients stratified according to their degree of epi-allelic shifting (as measured by EPM) between primary and recurring tumors.
D. Correlation between epi-allelic shifting during progression (as measured by EPM) and the time between first and second surgery. r: Pearson correlation.

**Supplementary Figure 7.**
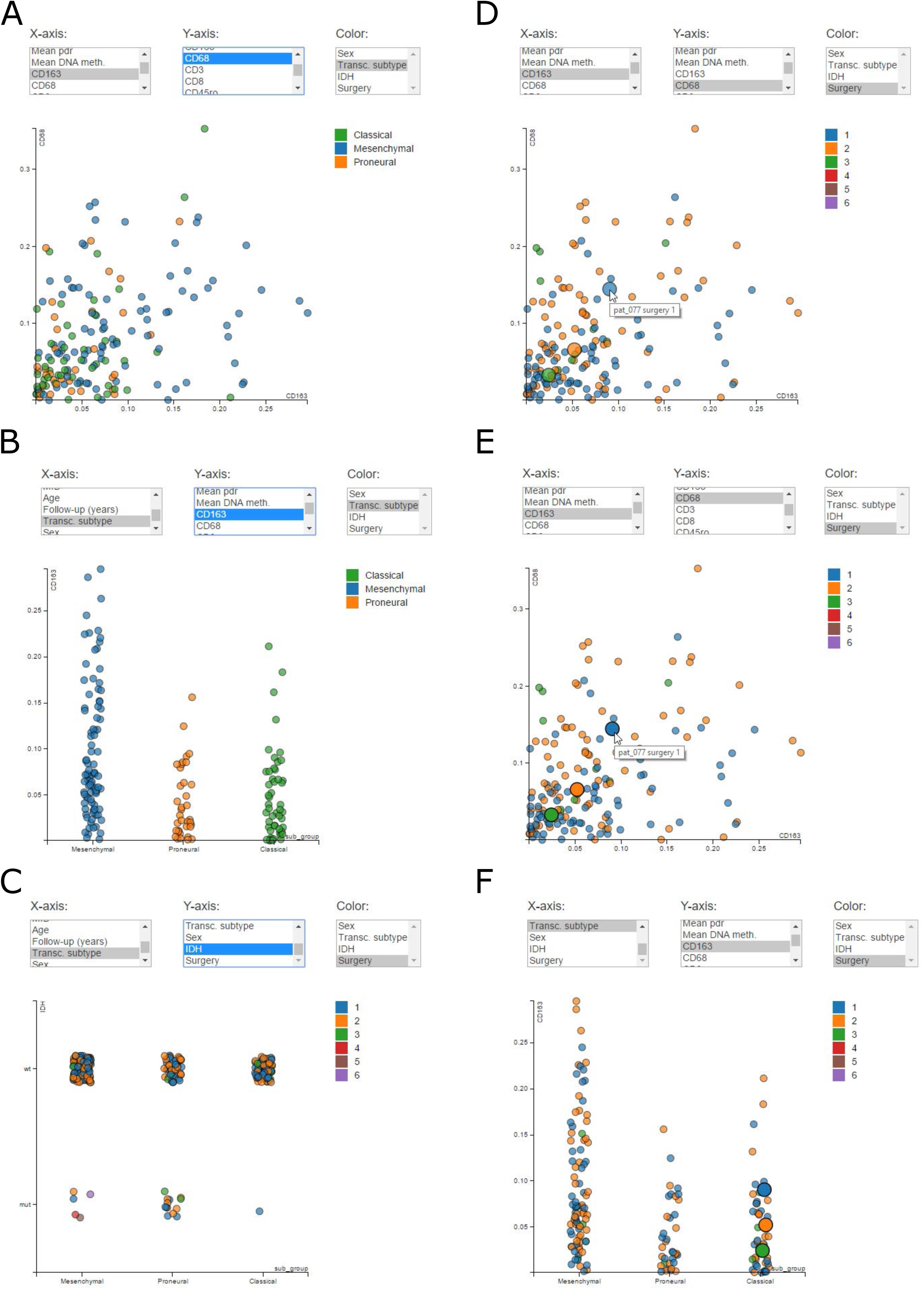
Illustration of the graphical data explorer on the Supplementary Website (http://glioblastoma-progression.computational-epigenetics.org/) A Comparison between two continuous variables.
B Comparison between one continuous and one categorical variable.
C Comparison between two categorical variables.
D Hovering over an individual data point shows information about the specific data point and also highlights all matched samples from the same patient.
E, F Clicking on a data point locks the highlighting (E) to follow the selected data point through additional analyses (F).

